# Bounding phenotype transition probabilities via conditional complexity

**DOI:** 10.1101/2024.12.18.629197

**Authors:** Kamal Dingle, Pascal Hagolani, Roland Zimm, Muhammad Umar, Samantha O’Sullivan, Ard A. Louis

## Abstract

By linking genetic sequences to phenotypic traits, genotype-phenotype maps represent a key layer in biological organisation. Their structure modulates the effects of genetic mutations which can contribute to shaping evolutionary outcomes. Recent work based on algorithmic information theory introduced an upper bound on the likelihood of a random genetic mutation causing a transition between two phenotypes, using only the conditional complexity between them. Here we evaluate how well this bound works for a range of genotype-phenotype maps, including a differential equation model for circadian rhythm, a matrix-multiplication model of gene regulatory networks, a developmental model of tooth morphologies for ringed seals, a polyomino-tile shape model of biological self-assembly, and the HP lattice protein model. By assessing three levels of predictive performance, we find that the bound provides meaningful estimates of phenotype transition probabilities across these complex systems. These results suggest that transition probabilities can be predicted to some degree directly from the phenotypes themselves, without needing detailed knowledge of the underlying genotype-phenotype map.

## I. INTRODUCTION

Mutation and selection are two key components of biological evolution. Ever since Darwin, there has been a preponderance of attention given to understanding how the latter (selection) operates [1, 2]. By contrast, understanding the processes of genetic mutation and how these mutations impact phenotypes and produce variation has historically received less attention [3]. More recently, the growth of evolution and development (evo-devo) is changing this imbalance [4], as is the study of genotype-phenotype maps [5, 6]. These fields are helping to uncover the quantitative laws underlying biological diversity by elucidating the mechanisms generating phenotypic variation [7, 8].

One important finding derived from these mechanistic approaches has been the realisation that upon random genetic mutation, certain phenotypes are more likely to appear than others, so that variation is not isotropic. In other words, many biological systems exhibit *developmental bias* [9–11] and *phenotype bias* [5, 6, 12–17], which both refer to the phenomenon that variation can be (strongly) biased towards certain phenotypic outcomes. Whether and in which ways these biases (and others [18–21]) in the introduction of variation might impact or steer evolutionary trajectories is a hot topic of research [20, 22–24]. For example, computer simulations have shown that when the bias is strong enough, it can lead to the fixation of sub-optimal phenotypes that are not the fittest, even while phenotypes with higher fitness values are available [17, 24–27]. Further, and perhaps more surprisingly, it has also been reported that the abundances of biomolecules in nature such as functional RNA and protein quaternary structures can be predicted using bias as a null-model [28–31]. These observations highlights the importance of studying bias in the arrival of variation.

A relatively recent addition to the analysis of biases was given in ref. [32], in which it was shown that in a general input-output map setting, under certain circumstances, randomly chosen inputs will lead to a bias for simpler, more regular or symmetric output patters, a phenomenon called *simplicity bias*. In particular, an upper-bound on the probability of outputs was presented, which was based on the complexity of the output shapes; in other words the probability *P* (*x*) that a certain shape or output *x* would appear from a random input can be bounded just using the complexity of the pattern, without requiring knowledge of the map details. This complexity-based bound was also applied to genotype-phenotype maps, which can be viewed as input-output maps. Specifically, it has been shown that protein quaternary structures, RNA secondary structures, gene-regulatory network concentration profiles, and biomorphs all exhibit simplicity bias [17, 30, 32]. These studies show that the information complexity of phenotype patterns can be a source of phenotype bias in biology. As an extension to the earlier theory of simplicity bias, Dingle et al. [33] derived a conditional form of the simplicity bias upper bound. This conditional version gives an upper bound on the probability *P* (*x→ y*) that some phenotype shape *x* transitions to some phenotype shape *y* upon a genetic point mutation to an underlying genotype. The bound is based on the conditional complexity of shape *y* given *x*, which measures how much extra information is required to make shape *y* given shape *x*. Because the precise value of transition probabilities *P* (*x → y*) depends on the details of genotype-space architecture, i.e. how exactly genotypes are assigned to phenotypes, it is not at all obvious that any predictions can be made regarding *P* (*x → y*) merely from the phenotype patterns *x* and *y* while completely ignoring the architecture. Hence, the discovery of an upper bound with non-trivial predictive success is noteworthy. Dingle et al. used their result to bound the probability of phenotype shape transitions for both RNA and protein secondary structures in computer simulations of the effects of mutations. Having said this, a significant observation is that for both of these molecular examples the genotype-phenotype connection is quite simple and direct. Therefore, it remains an important open question whether the bound on *P* (*x → y*) can also describe transition probabilities in less trivial and more realistic genotype-phenotype maps covering a wider range of biological phenomena.

In this work we seek to address this open question by expanding the investigation of the conditional form of the simplicity bias upper bound and testing its applicability in a range of genotype-phenotype maps. These maps are chosen specifically because they are either more complicated (in some sense), realistic, or have a less direct connection between genotypes and phenotypes. While our broader goal is to advance the understanding of genotype-phenotype map architecture in order to understand the processes and factors at play within biological evolution, here we will not perform evolutionary simulations of populations but instead restrict ourselves to the narrow biophysical question: upon a random mutation, can we predict (or bound) phenotype transition probabilities using conditional complexity? Answering this question can also be viewed as predicting the strength and direction of the genotype-phenotype map bias. The maps we study are: an ordinary differential equation circadian rhythm model; a matrix-multiplication model of gene regulatory networks; a model of tooth shape formation with different ways to measure complexity; a polyomino self-assembling model of protein quaternary shapes; and the well-known HP protein model. We assess three levels of transition probability prediction of increasing stringency. Our main findings are that the conditional simplicity bias appears in these more challenging maps (except perhaps the HP proteins), but rarely are all three levels of prediction are achieved, and we discuss possible reasons for these failures.

## II. BACKGROUND AND RELEVANT THEORY

### A. Genotypes, phenotypes, and bias

This study sits in the context of genotype-phenotype maps. In general, such maps refer to the association of genotypes to phenotypes, where genotypes refer to genome sequences, but also to some parameters which depend on gene sequences, while not necessarily being gene or nucleotide sequences themselves. Phenotypes are any traits, properties, and functions of an organism, and in general any properties that emerge reproducibly from the interaction of mostly hereditary generative factors, without stipulating that they are necessarily the final states of their generative dynamics. Examples include such things as eye colour, an animal’s height, and the mean number of eggs laid by a certain type of bird.

In this work we will be restricting our attention to a limited class of phenotypes, namely discrete shapes or patterns. Hence, the shape of a protein molecule is an example of a shape or pattern phenotype, but eye colour and height, for example, are not. The reason for this restriction is that we will be exploring the application of compression-based complexity to the probability of transitioning from one phenotype to another, and hence the phenotype must be such that a meaningful complexity can be obtained. Eye colour, and height, for example, do not admit meaningful compression-based complexity quantification. Of course, this restriction will limit the applicability of our work, but nonetheless there are still many maps which it potentially applies to.

Given some biological shape or pattern, which we take as a phenotype, the complexity of such a phenotype can be estimated via the complexity of the pattern. Taking an example from earlier work [33], an RNA secondary structure is routinely represented in bioinformatics as a “dot-bracket” string of characters, which denote the bonding pattern of the biomolecule. This string of characters defines a pattern, which has a meaningful complexity. Note also that the phenotype pattern must be repeatable (deterministic), in other words, the same genotype will yield the same phenotype. Contrast this with a hypothetical situation where a pair of genes randomly express, and time-varying environmental conditions also influence the phenotype pattern. In this scenario, the resulting pattern would likely be random and be different each time such a process is allowed to operate. While this type of random and complex pattern may be common in biology, we do not consider these and our body of theory would in all likelihood not apply.

The term *phenotype bias* refers to genotype-phenotype maps where the number of genotypes per phenotypes is (strongly) non-uniform [13, 34]. This concept is closely related to *developmental bias*, which refers to the fact that certain types of trait can be more readily made via the process of organismal development, or more generally to non-isotropic phenotype variation upon random mutations [11].

### B. AIT and algorithmic probability

Before looking at the applications of simplicity bias, we will briefly cover some of the related background theory. These details are given just for the sake of completeness, but will not (or only rarely) be directly invoked or used in this work.

Within theoretical computer science, *algorithmic information theory* [35–37] (AIT) connects computation, computability theory, and information theory. The central quantity of AIT is *Kolmogorov complexity, K*(*x*), which measures the complexity of an individual object *x* as the amount of information required to describe or generate *x. K*(*x*) is more technically defined as the length of a shortest program which runs on an optimal prefix *universal Turing machine* (UTM) [38], generates *x*, and halts. Intuitively, *K*(*x*) is a measure of the compressed version of a data object. Objects containing simple or repeating patterns like 010101010101 will have low complexity, while random objects lacking patterns will have high complexity.

An increasing number of studies show that AIT and Kolmogorov complexity can be successfully applied in the natural sciences, including thermodynamics [39–41], quantum physics [42], entropy estimation [43, 44], biology [30, 45, 46], other natural sciences [47], as well as engineering and other areas [48–50]. These applications typically use approximations to Kolmogorov complexity, which mostly take the form of real-world data compression algorithms (see more on this below). While real-world compression algorithms will fall short of accurately estimating the true Kolmogorov complexity, as these studies verify, the approximations often work very well.

An important result in AIT is Levin’s *coding theorem* [51], establishing a fundamental connection between *K*(*x*) and probability predictions. Mathematically, it states that

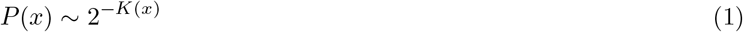

where *P* (*x*) is the probability that an output *x* is generated by a (prefix optimal) UTM fed with a random binary program. Probability estimates *P* (*x*) based on the Kolmogorov complexity of output patterns are called *algorithmic probability*. Given the broad-reaching and striking nature of this theorem, it is somewhat surprising that it is not more widely studied in the natural sciences. The reason in part for this inattention is that AIT results are often difficult to apply directly in real-world contexts, due to a number of issues including the fact that *K*(*x*) is formally uncomputable and the ubiquitous use of UTMs which may not be common in nature. See Appendix A for more discussion on applied AIT.

### C. Simplicity bias

Conscious of the difficulties associated with applying algorithmic probability in the natural sciences, approximations to algorithmic probability in real-world input-output maps have been developed, leading to the observation of a phenomenon called *simplicity bias* [32]. Simplicity bias is captured mathematically as

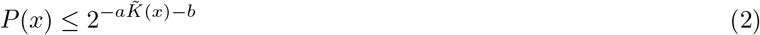

where *P* (*x*) is the (computable) probability of observing output *x* on a random choice of inputs, and is the approximate Kolmogorov complexity of the the output *x* which in general is some compression-based complexity measure, but the bound was also introduced and tested on other more general complexity metrics. See below for details on how we estimate complexity in this current work. In words, Eq. (2) says that complex outputs from input-output maps have lower probabilities, and high probability outputs are simpler. The constants *a >* 0 and *b* can be fit with little sampling and often even predicted without recourse to sampling [32]. See below for more discussion on the values of *a* and *b*.

Examples of systems exhibiting simplicity bias are by now wide ranging and include molecular shapes such as protein structures and RNA [30], finite state machines outputs [52], as well as models of financial market time series and differential equation systems [32], deep neural networks from machine learning [53–55], and dynamical systems [56, 57], among others. The ways in which simplicity bias differs from Levin’s coding theorem mentioned above include that it does not assume UTMs, uses approximations of Kolmogorov complexity, and for many outputs *P* (*x*) ≪ 2^−*K*(*x*)^. Hence the abundance of low complexity, low probability outputs [52, 58] is a signature of simplicity bias.

A full understanding of exactly which systems will and will not show simplicity bias is still lacking, but the phenomenon is expected to appear in a wide class of input-output maps under fairly general conditions. Some of these conditions were suggested in ref. [32], including (1) that the number of inputs should be much larger than the number of outputs, the number of outputs should be large, and (3) that the map should be ‘simple’ (technically of *O*(1) complexity) to prevent the map itself from dominating over inputs in defining output patterns. See Appendix B for more discussion on map complexity. Finally (4), because many AIT applications rely on approximations of Kolmogorov complexity via standard lossless compression algorithms [59, 60] (but see [61–63] for a fundamentally different approach), another condition proposed is that the map should not generate pseudo-random outputs like *π* = 3.1415… which standard compressors cannot handle effectively. The presence of such outputs may yield high probability outputs which appear ‘complex’ hence apparently violating simplicity bias, but which are in fact simple.

Note that it remains to be seen whether simplicity bias can appear in situations where one or more of these conditions is not met. For example, it has recently been shown that simplicity bias appears in the logistic map from chaos theory [56], a mapping known to be able to create pseudo-random patterns, hence potentially violating condition (4).

### D. Conditional simplicity bias

The upper bound in Eq. (2) is relevant when genotypes are randomly sampled from the full space of possible genotypes. However, in biology it is also important to consider phenotype transitions, that is, the probability that upon a genetic mutation the phenotype *x* becomes phenotype *y*. While single point mutations are perhaps the most common settings in mutagenesis experiments and evolutionary biology, studying situations where not just one but a few mutations are imposed is also interesting. With phenotype transitions in mind, in ref. [33] (but see also [64]) a conditional form of simplicity bias was derived, taking the form

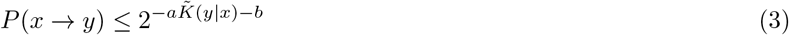

To elaborate, this bound says that the probability *P* (*x → y*) that phenotype *x* becomes phenotype *y* upon a point mutation (or a few) is modulated by the conditional complexity of *y* given *x*. It is necessary that the number of mutations be small, otherwise if large numbers of mutations are introduced, then it is effectively the same as a completely random genotype, for which Eq. (2) is more relevant. For *y* which are either simple or similar to *x*, then the probability *P* (*x → y*) can be high; for *y* which are complex and different to *x, P* (*x → y*) must be low.

Eq. (3) uses the conditional complexity, 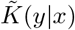, which measures the extra information required to generate some pattern *y*, given access to *x*. For example, if *x* = *abb* then the string *y* = *abbabbabb* can be generated by printing three repetitions of *x*, showing that we can use *x* to easily generate *y*, and hence 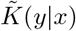 is small. Additionally, even if *x* is some complex pattern like *x* = *abaababbaaa* then the conditional complexity 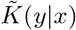 is small for *y* = *abaababbaaa* because we can generate *y* by simply copying *x*. On the other hand, if *x* and *y* share no similar patterns, then knowing *x* will not aid in generating *y* at all, and so 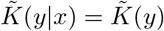. It follows that if either *y* itself is simple, or *y* is similar to *x*, then the conditional complexity will be low. If *y* is complex and different to *x*, then the conditional complexity will be high. To calculate the conditional complexity, the following relation [33] is used

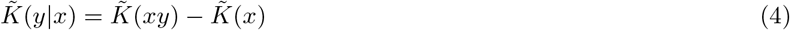

In this work we will primarily use the complexity measure based on Lempel-Ziv 1976 [59], denoted *C*_*LZ*_(*x*) as used earlier in multiple simplicity bias studies [32, 33, 52]. The authors Lempel and Ziv (and others) developed a handful of complexity estimators and compression algorithms, which became the basis for some data compression methods used today, such as gzip. There are variations in how each algorithm operates, but the common theme is that data objects are compressed into smaller sizes than their literal or raw forms, and these compressions are made possible by exploiting repeating patterns. Kolmogorov complexity equates “complexity” with incompressibility, and “simplicity” with compressibility, and so compression algorithms can be used as complexity measures.

### E. Application to genotype-phenotype maps

The conditional upper bound Eq. (3) can be seen as formalising and quantifying the intuition, perhaps held by biologists familiar with experimenting with genetic mutations, that mutations often do not have large effects [65], but occasionally rare mutations can have large effects. Going beyond a mere intuition however, the bound can be used to make predictions about which phenotypes are more or less likely to appear, and by how much [33].

It is noteworthy that we can make these non-trivial predictions, even while being agnostic about the details of the underlying genotype-phenotype map. To be truly agnostic about the map details, we should use phenotype complexity measures which do not assume or utilise any details of the genotype-phenotype mapping. In other words, ideally we should use complexity measures which utilise only the patterns of the phenotype if we are to make a theory for transition probabilities based only on phenotype shapes and patterns. Clearly, if the details of the genotype-phenotype map are known, or if data recording the effects of genetic mutations is available, then these can be used to make more accurate predictions of *P* (*x* → *y*) than by Eq. (3) alone. The primary use of Eq. (3) is to be able to make some non-trivial predictions even when such map details or data are not available.

## III. EXPERIMENT SET-UP

Before undertaking the numerical experiments, we will describe the general approach to be employed, and the different levels of prediction which will be assessed.

### A. Outline of methods and predictions

The protocol for the numerical experiments will be as follows: for each genotype-phenotype map, a suitable genotype and (discretised) phenotype will be defined. Then, via computational sampling and mutating genotypes, we will computationally estimate the transition probability *P* (*x → y*), representing the probability that the shape/pattern *y* appears as a phenotype when a random single point mutation is introduced to one random genotype underlying *x*. These transition probability estimates will be plotted against the upper bound in Eq. (3), and thereby test the accuracy of the bound.

### B. Approximating the complexity and constants in the bound

Eq. (2) has two parameters, *a >* 0 and *b*. Following refs. [32, 33], we take *b* = 0 as a default prediction, and we scale the complexity estimates aiming to make *a* ≈ 1, using the following expression

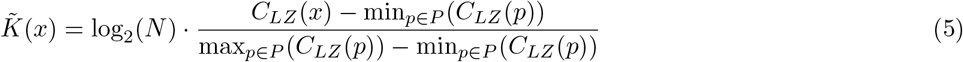

where *N* is the total number of phenotypes, *P* is the set of all phenotypes, and the max and min are taken over all *N* phenotypes. The purpose of the scaling is to make 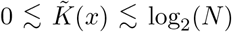, which is the typical range of Kolmogorov complexity for *N* different patterns (with some assumptions [49]). For example, if the set of *N* patterns consists of all binary strings of length *L*, then there will be *N* = 2^*L*^ strings, with complexity ranging up to ∼log_2_(*N*) = *L* bits.

This particular scaling of Eq. (5) is not fundamental to the theory of simplicity bias, and is used mainly because the complexity measure *C*_*LZ*_ returns values which (presumably) correctly order the output/phenotype patterns in terms of complexity, but gives overly large absolute complexity values (a problem which is especially relevant for short strings). Other scaling methods could be used, and indeed in ref. [32] a slightly different scaling was used that did not contain the minimum term. An exception in this work regarding this scaling is when studying teeth complexity, where we use a different approach since our complexity measurement relies on specific tooth traits (below) rather than the *C*_*LZ*_ complexity measure that is used throughout the rest of the paper.

For the conditional complexity, 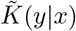, the same complexity scaling method is used as in Eq. (5), but instead of the term log_2_(*N*), log_2_(*N*_*y*_(*x*)) is used, where *N*_*y*_(*x*) is the number of different phenotypes *y* such that *P* (*x* → *y*) *>* 0, i.e., the number of accessible phenotypes via mutation from *x*. Similarly, the max/min is taken over the set of accessible phenotypes via mutation from *x*. It is possible that the conditional complexity scaling might vary with *x*, because different starting phenotypes might have different numbers *N*_*y*_(*x*) of accessible phenotypes in its neighbourhood. If so, it will affect the scaling only, and not the ordering of the complexities of the phenotypes.

In some genotype-phenotype maps estimating *N* or *N*_*y*_(*x*) is possible a priori. For example, if it is known that the set of accessible phenotypes corresponds to the set of all possible binary strings of length *L*, then this implies the number is 2^*L*^. In many cases, the set of possible accessible phenotypes is not known, or cannot easily be estimated. With this scaling of Eq. (5), we expect *a* ≈ 1 to model the data well, but in practice this may not occur. This error may be because an accurate estimate of *N*, or of the max/min values, is not possible to obtain. It can also occur when the output (phenotype) is corrupted by random noise [57] instead of being deterministically encoded by the input/genotype. Alternatively, it may be that there is not enough bias in the map (i.e., the probabilities are too uniform), and hence the bound of Eq. (2) or Eq. (3) is not followed closely. If the default predictions of *a* = 1 and *b* = 0 are not observed to be accurate, then these could be obtained (if needed) by fitting to the upper bound, that is, the highest probability value for each unique complexity value.

These requirements for the scaling in Eq. (5) represent a central challenge in predicting the quantitive slope of the upper bound decay, and hence a main challenge is making a priori transition probability predictions. It follows that we may observe conditional simplicity bias in a general loose sense, but without a precise correct prediction for the upper bound slope. It is worth highlighting that, as explored in ref. [33] even without good estimates of *a* and *b* it is still possible to make non-trivial and useful predictions, such as which of two possible phenotypes *y*_1_ and *y*_2_ is more or less likely to appear upon random mutation. The reason this is possible is that according to the bound, constants *a* and *b* are not required to infer which phenotype is more or less likely, only to make an estimate of the actual probability value. Relatedly, if *a* can be estimated but *b* cannot, then it is still possible to estimate the relative probabilities because the value of *P* (*x → y*_1_)*/P* (*x → y*_2_) does not depend on *b*.

### C. Levels of conditional simplicity bias

In terms of predictions deriving from conditional simplicity bias theory discussed above, we are interested in three closely related but nonetheless non-equivalent phenomena. We will call these three *levels* of conditional simplicity bias:

**Level (I)**: log_10_ *P* (*x*→ *y*) tends to decrease with increasing conditional complexity 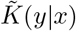, but not necessarily with a linear upper bound.

**Level (II)**: log_10_ *P* (*x*→ *y*) tends to decrease with increasing conditional complexity 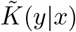, with a linear upper bound, but not necessarily with the predicted slope.

**Level (III)**: log_10_ *P* (*x*→ *y*) tends to decrease with increasing conditional complexity 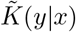, with a linear upper bound, including correctly predicting the slope.

These three situations are increasingly precise and stringent. If any of these levels are apparent in the example genotype-phenotype maps, we will consider it a predictive success for the theory. The slope especially is difficult to estimate (as discussed above), hence achieving Level (III) will be a challenge. In the earlier study of ref. [33] looking at RNA and protein secondary structure, Level (I) and Level (II) were achieved in all cases, while Level (III) was achieved for only some test-case phenotypes.

To test the statistical significance of the three levels, we do the following tests: For Level (I) we use Spearman’s rank correlation coefficient *ρ* using all the data points available in each test case (not just the upper bound). A significant negative *ρ* indicates that Level (I) is achieved. We take *p* − *value <* 0.05 as the threshold of significance. For Level (II) we use Pearson’s *R*^2^ to measure the degree of linearity of the upper bound of the relationship between conditional complexity and transition probability for each test case. In this context, *R*^2^ represents the proportion of the variance in the transition probability that can be explained by the conditional complexity using a linear model. High values of *R*^2^ indicate a strong linear relationship. Note that by ‘upper bound’ we mean the highest log_10_ *P* (*x* → *y*) value for each unique conditional complexity value 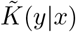. We consider that Level II has been achieved only if *R*^2^ *>* 0.5.

For Level (III), we use a bootstrap method to determine if two slopes are significantly similar by generating a distribution of slope differences. Starting with the slope given by the bound model, we independently resample the observed maximum probability values for each conditional complexity category of the test case, with replacement, to create 1000 bootstrap samples. Importantly, we do not recalculate new upper bounds for each bootstrap sample; we resample from the existing upper bound points. For each sample, we calculate a slope and then subtract the bound model slope from it, producing a distribution of differences. By calculating the 95% confidence interval (CI) of these differences, we assess if 0 lies within the interval. If 0 is inside the CI, it suggests that the slopes are not significantly different, indicating they are similar; if 0 is outside the CI, the slopes are significantly different. To apply this method for assessing whether the predicted slope is significantly different from the upper bound slope, we first verify that the upper bound model meets both of the following requirements: *R*^2^ *>* 0.5 and the data can be significantly explained by a linear model (*p* − *value <* 0.05).

It should be kept in mind that the metrics used to measure Level (I), (II), and (III) have limitations, and are only roughly indicative of the success of the conditional simplicity bias predictions. By this we mean that a high Spearman’s correlation value or a high *R*^2^ value, for example, do not imply that Level (I) or (II) are convincingly achieved. On the other hand, we need *some* kind of metrics to assess the prediction performance, so these will suffice.

## IV. NUMERICAL EXPERIMENTS

### A. Choice of genotype-phenotype maps

We now proceed to a number of numerical experiments, testing the conditional simplicity bias bound Eq. (3) for a variety of different maps which are summarize in Table I. As discussed above, the conditional simplicity bias under investigation is intended to apply to a certain restricted class of genotype-phenotype maps. More specifically, we will be focussing on maps for which there are: many more genotypes than phenotype (redundancy); a strongly non-uniform distribution of genotype to phenotypes (phenotype bias); a finite collection of phenotypes which are either shapes or patterns, and for which some meaningful compression-based complexity values can be measured. In addition to testing a wider range of genotype-phenotype maps in this current study, where appropriate, we also test some other complexity measures.

**TABLE I.**
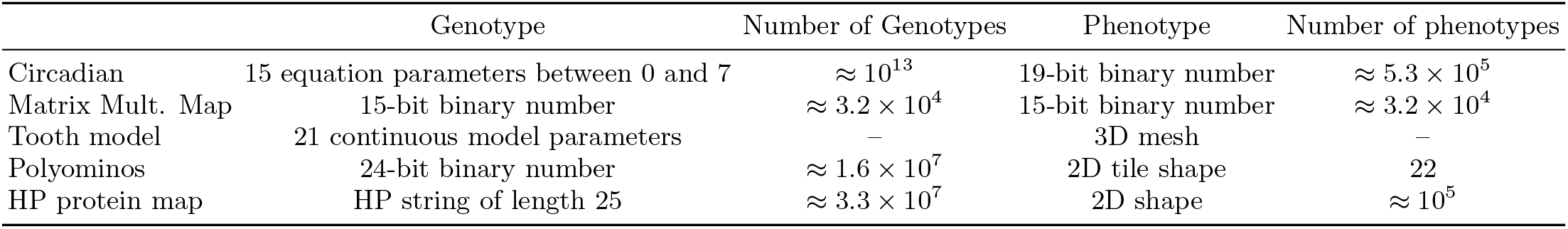
Summary of models. The figure summarizes distinct genotype and phenotype classifications alongside their corresponding numerical possibilities within each biological model.

|  | Genotype | Number of Genotypes | Phenotype | Number of phenotypes |
| --- | --- | --- | --- | --- |
| Circadian | 15 equation parameters between 0 and 7 | $\approx 10^{13}$ | 19-bit binary number | $\approx 5.3 \times 10^5$ |
| Matrix Mult. Map | 15-bit binary number | $\approx 3.2 \times 10^4$ | 15-bit binary number | $\approx 3.2 \times 10^4$ |
| Tooth model | 21 continuous model parameters | — | 3D mesh | — |
| Polyominos | 24-bit binary number | $\approx 1.6 \times 10^7$ | 2D tile shape | 22 |
| HP protein map | HP string of length 25 | $\approx 3.3 \times 10^7$ | 2D shape | $\approx 10^5$ |

### B. Circadian rhythm

For our first genotype-phenotype map, we will study transition probabilities of a discretised circadian rhythm differential equation model, developed by Vilar et al. [66]. The model was originally introduced to study strategies that biological systems use to minimize noise in circadian clocks, but was also studied in ref. [32] in the context of simplicity bias, and we follow their definitions of inputs (genotypes) and outputs (phenotypes). In this model, ‘genotypes’ are the values of the 15 equation parameters, which are defined as integers between 0 and 7. Hence there are 7^15^ ∼ 10^13^ possible genotypes.

As for the resulting phenotype *x* associated to each parameter-combination genotype in this model, again following ref. [32] we take the chemical time-concentration profile of the activator, as defined in [66]. This profile has a certain shape which depends on the genotype parameters, and the shape denotes the concentration level of a product at the end of the regulatory cascade over time. To define the phenotype profile *x*, we discretise the shape into an “up-down” fashion [67, 68]. For this, every 25000 regular time intervals the slope of the curve is computed, and we write “1” if the slope is positive, and “0” if it is negative or flat. Since we model the circadian cycle for 499,999 iterations, we obtain 19 intervals, each of them defined by a “0” or “1”, these gives us the phenotype of a specific circadian cycle. The reason for discretising is that calculating both the complexity and probability of continuous curves is problematic, and simplicity bias theory has been developed in the context of discrete output shapes/patterns, which are typically binary strings. In this manner, we have framed the genotype-phenotype map as a map from 15 parameters to length 19 binary strings. The standard *C*_*LZ*_ complexity measure will be used for this map, and scaled according to Eq. (5). The choice of phenotype length to be 19 comes as a trade-off between not being too short such that only very few phenotypes are possible, and not being too long such that so many phenotypes are possible that obtaining decent statistics from frequency sampling becomes overly taxing.

It will be apparent that in this circadian rhythm model, the connection between the input parameter genotypes and output shapes is non-trivial and also not at all direct. Hence this map will act as a good test-case for the applicability of Eq. (3).

To test the predictive capacity of Eq. (3), we need to have good estimates of the true value of *P* (*x* → *y*), for some phenotype *x*, to compare against. In principle, the exact value of *P* (*x*→ *y*) could be found via complete enumeration of all genotypes and computing all possible single point mutations, but typically that approach is computationally very demanding. Further, to compute *P* (*x* → *y*) for all possible pairs of *x* and *y* is even more demanding, so in practice we will choose only a few test-case phenotypes *x*, and then undertake uniform random sampling of the neutral space of *x* (i.e., the set of genotypes associated to *x*). The test-cases are obtained from sampling 100,000 random genotypes. These samples were randomly chosen while imposing the condition that the test-cases should have a spread of complexity and/or probability values. The reason for imposing this spread is to test the bound of Eq. (3) in a variety of complexity/and or probability cases. If instead we simply took the first few phenotypes to appear on random genotype sampling, then potentially the only phenotypes to appear would be very high probability, and therefore likely simple, or have some other narrow set of properties not typical of the full space of phenotypes.

For each of the collection of (*n* = 35) test-case starting phenotypes *x*, we found the genotypes that give rise to the same phenotype in the original 100,000 sample, that is, we find all the genotypes belonging to the same neutral set that exist within the 100,000 sample. For each of these *x*, we mutated each sequence with all possible single point mutations. In the case of the circadian rhythm, there were 15 parameters with each taking 8 possible values so that there are 7 possible mutations for each parameter. Therefore, for each sequence there are 15 *×* 7 = 105 possible resulting sequences after all possible single point mutations are imposed. Note that these resulting sequences will not necessarily be distinct. For each of these 105 resulting sequences, we can use the genotype-phenotype map to find their resulting phenotype *y*. By counting the frequency with which each *x* transitions to each of these *y* phenotypes, computational estimates of *P* (*x* → *y*) can be found.

Figure 1 shows the results of the above-described computational experiments. In panel (a), the basic probability-complexity plot is displayed for completeness. As can be seen, there is a roughly linear upper bound decay in log *P* (*x*) with increasing complexity 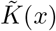, as in Eq. (2). In (b), (c), and (d) the conditional simplicity bias data is plotted for three test-case starting phenotypes *x*. In Figure 1, and subsequent figures, the examples illustrate varying degrees of success in predicting simplicity bias levels. The progression starts with cases where all levels are successfully predicted, followed by examples with fewer levels predicted, when such examples exist. Within each category of success, the examples were selected randomly. To see all the cases we explored see supplementary information. As is apparent in (b), there is a decay in transition probability *P* (*x* → *y*) with increasing conditional complexity, as can be seen by their negative correlation, fulfilling the requirements for Level (I). Additionally, in these examples, (b) and (c) show a *R*^2^ *>* 0.5, which indicates a linear correlation of the upper bound. However (d) fails to fulfill the requirements of Level (II). Finally, comparing the slopes of the bound model (in red) and the slopes in the 95% CI of the upper bound data point, calculated using bootstrapping with replacement, we determined if the slopes are significantly different or if the bound model correctly predicted the slope of the upper bound decay. In the examples shown here, only (b) correctly predicts the slope. The plots show a fitted upper bound, in black, and also an upper bound prediction, in red, assuming Eq. (5) works for the bound slope.

**FIG. 1.**
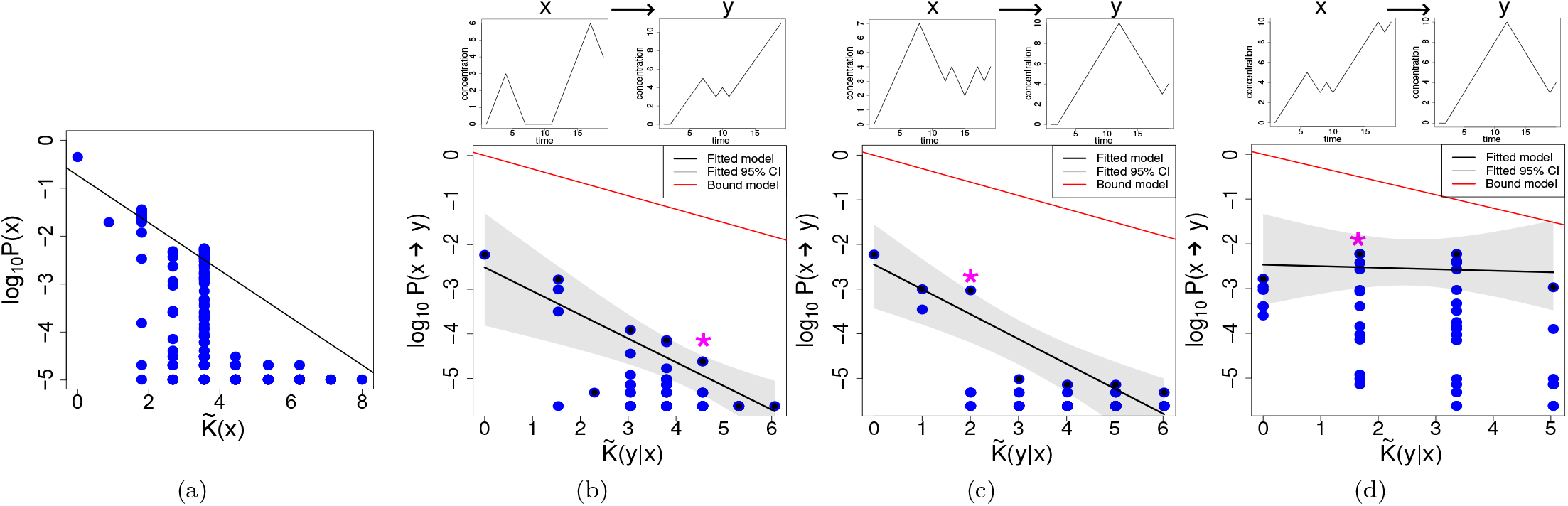
Circadian rhythm map. (a) Simplicity bias observed for uniform random sampling the circadian rhythm map genotype sequences. The black line shows the linear regression using the maximal values for each complexity value, 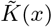. The following examples illustrate cases where the three levels of simplicity bias succeed to varying degrees. On top of each plot we can see the phenotype x on the left side and an example of a phenotype y, into which phenotype x has mutated to. On the plots, the * indicates where the example y can be found. The following three examples are chosen to represent different levels of success for the different levels of simplicity bias, whenever such an example exists, (b) shows an example when all levels are achieved, (c) when only the I and II levels are achieved and (d) when only the I level is achieved. This approach to choosing display figures will be used for all the maps we study. (b) Example for the transition probabilities *P* (*x* → *y*) of a starting phenotype with complexity 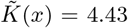. Each blue point shows the conditional complexity 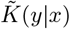) of one of the mutants found in the 1-mutant neighborhood exploration. The black line and gray shaded area show the linear regression performed using a bootstrap approach as described in the main text. The red line shows the bound model calculated using Eq. (3). In this example Spearman’s correlation *ρ* =− 0.44, such a negative correlation confirms Level (I). Level (II) is also achieved, because *R*^2^ = 0.71. Finally, Level (III) is also achieved, because the differences of the slopes calculated with the bootstrap method and the slope of the bound model are not significantly different from 0. (c) In this example, using a starting phenotype of complexity 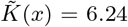, Level (I) is achieved with a *ρ* = − 0.34. Level (II) is also achieved, having a *R*^2^ = 0.85, however, Level (III) is not attained, because slopes of the bound and fitted model are significantly different. (d) In this example, using a phenotype of complexity 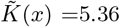, Level (I) is achieved with a *ρ* = − 0.47, even though the negative relation is very weak. However, Level (II) is not achieved, since *R*^2^ = 0.04 is below the 0.5 threshold we established. Level (III) is not achieved since the slopes are significantly different.

The results shown in Table II summarize the data by calculating the average and variability for each phenotype, considering all the genotypes that were found. When calculating the averages, the results from each genotype linked to the phenotype are equally included. Thus, if multiple genotypes were found for a phenotype, each contributes to the final calculation of averages and standard deviations, reflecting the expected outcome if one were to randomly select a genotype for a given phenotype. In addition, in Table III, we show similar results, but focusing on the average and variation of the phenotypes, without considering how many genotypes map to each of them. Level (I) is achieved, with all cases showing a negative correlation. Similarly, Level (II) is achived in many cases showing a *R*^2^ *>* 0.5. Level (III) shows a lower degree of success, with only 14% of the slopes being predicted.

**TABLE II.** Statistics for Levels of simplicity bias. **Genotype**. For each of the models we researched, we present the average and standard deviation that a random genotype will have at each level of simplicity bias. Additionally, we show the proportion of prediction for each level. The proportions are based on the total number of genotypes predicted in the previous level.

|  | Level I |  | Level II |  | Level III |
| --- | --- | --- | --- | --- | --- |
| | $\rho$ | Pred | $R^2$ | Pred | Pred |
| Circadian | $-0.47 \pm 0.01$ | 0.99 | $0.60 \pm 0.01$ | 0.99 | 0.14 |
| Matrix Mult. Map | $-0.21 \pm 0.01$ | 0.89 | $0.78 \pm 0.01$ | 0.52 | 0.67 |
| Tooth Cusps | $-0.84 \pm 0.07$ | 0.92 | $0.87 \pm 0.01$ | 1.00 | 0.91 |
| Tooth OPC | $-0.53 \pm 0.05$ | 0.92 | $0.63 \pm 0.01$ | 0.84 | 0.66 |
| Polyomins | $-0.43 \pm 0.01$ | 0.53 | $0.69 \pm 0.01$ | 0.52 | 0.06 |
| HP protein map | $-0.71 \pm 0.02$ | 0.49 | $0.77 \pm 0.01$ | 0.47 | 0.95 |

**TABLE III.** Statistics for Levels of simplicity bias **Phenotype**. For each of the models we researched, we present the average and standard deviation that a random genotype will have at each level of simplicity bias. Additionally, we show the proportion of prediction for each level. The proportions are based on the total number of genotypes predicted in the previous level.

|  | Level I |  | Level II |  | Level III |
| --- | --- | --- | --- | --- | --- |
| | $\rho$ | Pred | $R^2$ | Pred | Pred |
| Circadian | $-0.47 \pm 0.01$ | 0.95 | $0.69 \pm 0.01$ | 0.79 | 0.63 |
| Matrix Mult. Map | $-0.21 \pm 0.01$ | 0.88 | $0.77 \pm 0.01$ | 0.54 | 0.68 |
| Tooth Cusps | $-0.84 \pm 0.08$ | 0.92 | $0.87 \pm 0.01$ | 1.00 | 0.91 |
| Tooth OPC | $-0.52 \pm 0.06$ | 0.92 | $0.63 \pm 0.01$ | 0.83 | 0.67 |
| Polyomins | $-0.60 \pm 0.01$ | 0.39 | $0.69 \pm 0.02$ | 0.28 | 0.28 |
| HP protein map | $-0.75 \pm 0.02$ | 0.49 | $0.78 \pm 0.01$ | 0.47 | 0.94 |

It should also be noted that we expect that point mutations on *x* will be most likely to produce phenotypes *y* which are similar to *x*, or have lower conditional complexity. Conversely, we might expect mutations to rarely produce phenotypes *y* with high conditional complexity values. This expectation is due to the conditional complexity bias bound assigning higher probabilities to phenotypes with similar or lower conditional complexity, effectively excluding those with higher conditional complexity. We tested this conjecture (Appendix B) and observed that indeed, in all studied cases, more frequently found phenotypes consistently show lower conditional complexity.

### C. Gene regulation network vector-matrix map

For our second genotype-phenotype map, we will use the vector-matrix multiplication map, which has been used to model genotype-phenotype maps [13, 32]. The map is defined by the following equation

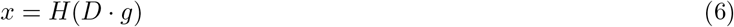

where *g* is a length 15 binary string genotype, *D* is a square matrix made up of randomly chosen entries taking values in *{*− 1, 0, 1}. These values can e.g. represent gene regulation, including promoting and suppressing gene expressions levels. Finally, the Heaviside function *H* is applied so that for component *j* of the vector *D* · *g* if the value is positive, then the *j*th component of *x* is set to 1, otherwise it is set to 0. In this manner, we have a map from binary strings of length 15 to binary strings also of length 15. Due to the action of the Heaviside function, there are typically many genotypes per phenotype.

This map is very abstract, and does not specifically model any one biological genotype-phenotype map, but was instead introduced to simulate in an abstract way the ‘computation’ of a phenotype pattern from a genotype. One interpretation of the binary string phenotype could be merely the presence or absence of a list of traits, e.g., the organism has/does not have blue eyes, or has/does not have wings. In this case, the order of the phenotype binary string would have no relevance, and hence the complexity of the binary sting would not be meaningful. However, given the abstract and general nature of the map, the binary phenotype could also be understood to be some biological pattern, structure, or shape. In this case, the complexity would be meaningful. We will assume this second case in what follows.

While the connection between genotype and phenotypes is quite direct and simple in this map, it is an interesting map because in ref. [32] it was shown that this map does *not* show simplicity bias. The reason as discussed in [32], is that the map itself has high information content: the matrix *D* contains *L*^2^ random values for a genotype of length *L*. Therefore the information content of the map itself is typically much higher than that of any genotype (*L*^2^ ≫ *L*). One of the conditions proposed for observing simplicity bias was to have a simple (technically *O*(1) complexity) map, which does not hold in this case. Even though it does not exhibit simplicity bias when sampling over the full range of inputs (genotypes), it is interesting to see if there is a kind of conditional simplicity bias in the transitions *P* (*x → y*).

In Figure 2(a) we see, as shown already earlier [32], that there is no simplicity bias with this map. Although there does appear to be some kind of *positive* relation of complexity and probability in this plot, this is due to the fact that there are many more higher-complexity binary strings, so there is greater chance of at least some of them having higher probability. Looking to Figure 2(b), (c), and (d), we still see that Level (I) is achieved, as also shown in Table II, in most cases we see that there is conditional simplicity bias, with the fitted upper bound decaying with increasing conditional complexity. Because the median correlation value is only about -0.18, the relation is not very strong. The upper bound decay is often significantly linear, as seen in Table II, therefore achieving Level (II), although there is a wide spread of cases which cannot be considered to decay in a linear way. However, only in 30% of cases is the upper bound model correctly predicted. It appears that the upper bound slopes, while roughly linear, do not have very steep slopes. In ref. [57] it was observed that the slope of the decay in the upper bound can be reduced when random noise is introduced to the outputs. Speculating, it may be that the slopes here are not steep due to the high complexity (hence randomness) of the map itself.

**FIG. 2.**
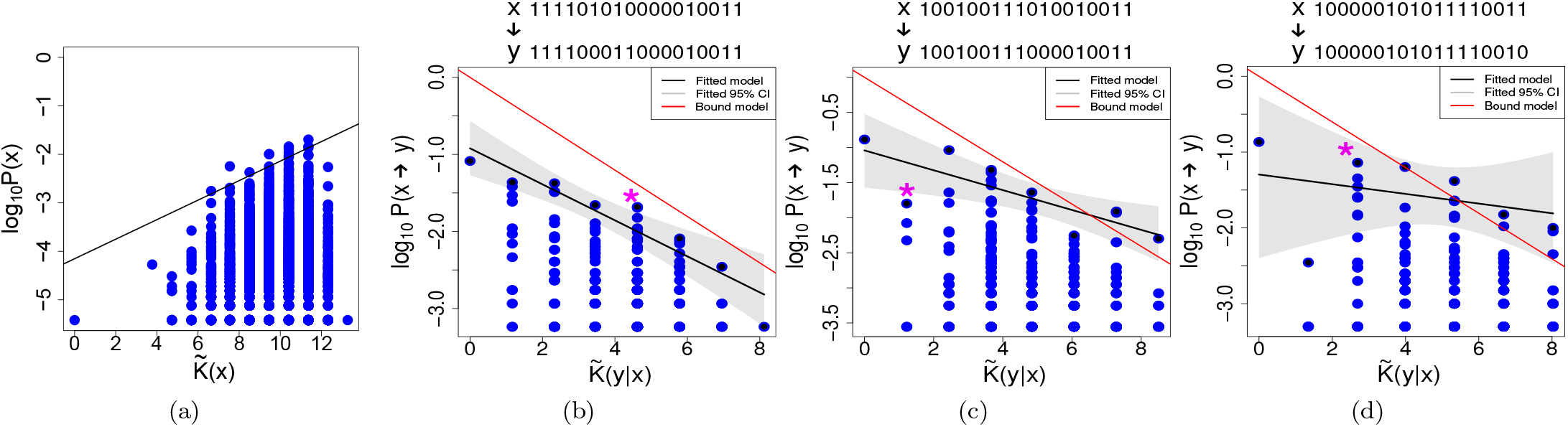
Matrix multiplication map. (a) No simplicity bias observed for the uniform random sampling of the multiplication matrix map. As before, the following examples illustrate cases where the three levels of simplicity bias succeed to varying degrees. On top of each plot we can see the phenotype x on the left side and an example of a phenotype y, into which phenotype x has mutated to. On the plots, the * indicates where the example y can be found. As previously explained, the following examples illustrate varying levels of success corresponding to different degrees of simplicity bias. (b) Example for the transition probabilities *P* (*x* → *y*) of a phenotype with complexity 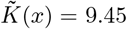. In this example *ρ* = − 0.26, this negative correlation confirms Level (I). Level (II) is also achieved, with a *R*^2^ = 0.89. Finally, Level (III) is also achieved, since the differences of the slopes calculated with the bootstrap method and the slope of the bound model are not significantly different from 0. (c) In this example, using a phenotype of complexity 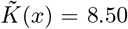, Level (I) is achieved with a *ρ* = − 0.24. Level (II) is also achieved, with a *R*^2^ = 0.64, Level (III) is not predicted, as slopes of the bound and fitted model are significantly different (d) In this example, using a phenotype of complexity 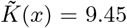, Level (I) is achieved with a *ρ* = − 0.22. However, Level (II) is not achieved, since *R*^2^ = 0.10, failing at this Level (II) also means that Level (III) is not achieved.

How is it possible that we observe conditional simplicity bias, but not simplicity bias when sampling purely random genotypes? The reason is possibly related to the following: If a map assigns genotypes to phenotypes in a purely random manner, then this is a maximally complex map. We can estimate the Kolmogorov complexity of the random map by taking the logarithm of the total number of possible ways to assign *n*_*g*_ genotypes to *n*_*p*_ phenotypes. For this computation, we can use Stirling numbers of the second kind, denoted *S*(*n*_*g*_, *n*_*p*_), and multiply by (*n*_*p*_!). For large *n*_*g*_ and *n*_*g*_ ≫ *n*_*p*_, which is typical in genotype-phenotype maps, then it is simpler to approximate that the number of maps is 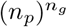. Hence the complexity of a random assignment of genotypes to phenotypes is roughly *n*_*g*_ log_2_(*n*_*p*_) bits. On the other hand, the complexity of the matrix map can be estimated as *O*(*L*^2^) bits, because there are *L*^2^ entries in the matrix, where *L* = log_2_(*n*_*g*_) is the length of the binary genotype. Clearly, *n*_*g*_ log_2_(*n*_*p*_) ≫ *O*(*L*^2^), and hence the map is far from being completely random. It follows that the mapping is of medium complexity, we could say: it has higher complexity than any one genotype and so is complex enough to impact the probability-complexity connection, but at the same time contains much less information than a purely random map (for which we would expect no connection between probability and complexity). Due to the non-random nature of the matrix map, we might expect to see some structure and pattern in how inputs are assigned to outputs. Hence this could explain the observation of conditional simplicity bias, even without the original form of plain simplicity bias.

### D. Tooth developmental model

Development of complex organs typically encompasses sets of different generative factors that can interact in non-linear ways [69]. For instance, genes may interact with each other dynamically to orchestrate changes in cells and tissues in a spatially and temporally distinct manner, while bio-mechanical parameters may bias which specific shape changes are possible or facilitated [70, 71]. Thus, numerical models exploring such developmental dynamics often feature heterogeneous input variables that transform simple patterns into complex ones whose values may be continuous and non-finite. This means that the emerging genotype-phenotype maps differ from many of the previously studied ones, namely regarding both non-discreteness of input and output variables as well as the heterogeneity of the mechanics of their interactions. It is therefore an interesting and important task to assess to what extent the mathematical laws established through the study of simpler models apply to this class of models too.

A representative numerical model of tooth developmental is in ref. [72], which allows testing of the contribution of genetic, cellular, and mechanical factors to the formation of realistic tooth shapes as folded 3D meshes. This tool has been used and modified throughout a number of studies in different organisms, namely rodents, seals, prehistoric mammals and, recently, sharks [72–76], underpinning its versatility and scientific pertinence. Besides testing mechanistic developmental hypotheses, this model allows for the study of trait evolution by parameter mutations [77]. It has also been a useful tool to explore genotype-phenotype map properties, revealing a bias against complex shapes [78] and morphospace degeneracy [75]. Here we build on this work by systematically assessing whether we observe comparable phenotype transition probabilities as with the previous models. We take advantage of the versatility of this tooth model by applying the model to tooth shapes of its original species of study, the seal *Pusa hispida* [72]. While the range of shapes in our (and the original) analysis recapitulates dental variation in a seal species [72], we emphasize that the model does not claim to be specific to this mammalian clade, but is capable of producing tooth shapes reminiscent of many other taxa by adjusting the ranges of developmental parameters [73, 74, 76, 79–82]. In this study, we quantify phenotypic complexity in two complementary ways, thus testing the generality of our hypotheses.

The first way to quantify tooth complexity is simply counting the number of cusps in a tooth [73, 83]. Here, we define cusps as local elevations on the *in silico* mesh representing the epithelial-mesenchymal interface, with local elevations identified as mesh nodes whose z-coordinate exceeds that of their neighbours.

A second way to measure tooth complexity is using Orientation Patch Count Rotated (OPCR), a widely-used, high-resolution metric for quantifying the surface complexity of teeth [76, 84–87]. A patch is defined as a group of contiguous points on the tooth surface facing the same “compass” direction, such that they have similarly angled normal vectors when projected on the XY plane [84, 86]. Orientation Patch Count (OPC) counts these distinct patches and approximates the number of ‘tools’ on the tooth crown used for breaking down food [84]. OPC has been shown to correlate with diet [84, 87, 88], with dental complexity increasing from hypercarnivory through omnivory to herbivory [84, 85]. This increase in surface complexity may reflect the increased demands of mechanical processing in herbivore diets, compared to that of carnivores.

Patch count provides a more sensitive measure of complexity compared to landmark-based methods due to its finer resolution of surface data [86]. OPC has been employed to measure the surface complexity of teeth in primates [86, 87, 89, 90], multituberculates [91], carnivorans [84, 85], rodents [73, 76, 84, 85], bats [88], and generalized models of tooth development and adaptation [78, 81]. OPCR further improves upon OPC by reducing sensitivity to tooth orientation [85, 86]. Using MorphoTester, a GIS software, we divide the tooth surfaces into patches of equivalent orientation, with a minimum patch size of three grid points [86]. We rotate individual molar specimens eight times across a total arc of 45° (5.625° per rotation), calculating OPC at each rotation, and averaging these eight values to obtain OPCR [86]. OPCR can then be visualized by colouring surface patches one of eight colours corresponding to patch orientation [86].

To accommodate the continuous map inputs, we first define our inputs as the unique combinations of 26 parameters responsible for cellular and genetic interactions in seal tooth development [72]. We then establish a biologically realistic range for each variable parameter by individually varying parameters until the tooth produced either an unrealistically flat structure or unrealistic globular clusters of cusps. Using these ranges, we apply Latin hypercube sampling to divide each range into 19,000 equal-probability strata, selecting one sample from each stratum. For the conditional simplicity bias experiments, we begin with a range of discretized genotypes (parameter combinations) that replicate real seal teeth found in nature [72]. Mutations are then introduced by modifications of the model parameters. Specifically, for each mutant, we changed a randomly chosen parameter *p* to a value ranging between the value of the respective parameter in the “parental” tooth *p*_0_ and either the minimal or the maximal allowed parameter value *p*_*m*_ (as in [75]). Since we use continuous values in this model, we could not explore every single possible mutation. Instead we explored 19,000 mutants per “parental” tooth. We calculated each mutant’s conditional complexity as the smaller value between the mutant’s OPCR and the absolute difference in OPCR between the parent and mutant tooth.

The results for the tooth model using cusp number as the complexity measure can be seen in Figure 3. In Figure 3(a) we show that despite the added complexity of the tooth model genotype-phenotype map, the probability of finding simple teeth is consistently larger than of finding more complex teeth, which is expectedly consistent with other models. In fact, the decrease of the logarithm of frequency with increasing phenotypic complexity follows a near-linear curve, with the notable exception of mono-cuspid teeth, the lowest possible complexity. The high frequency of very simple shapes (1 cusp) might reflect that this minimum complexity exists for free, i.e., without the activation of specific mechanisms, and can be equally accessed from any part of the morphospace. As shown in Figure 3(b-d), we see that Level (I) is achieved in these examples. In 3(b-c) Level (II) and (III) are also achieved, showing that the decay is linear and the slope of decay can be predicted. However, in 3(d) although Level (II) is achieved, Level (III) is not successful. In Table II we can see that in all cases Level (I) and (II) is achieved, but Level (III) is successful in 57% of the cases.

**FIG. 3.**
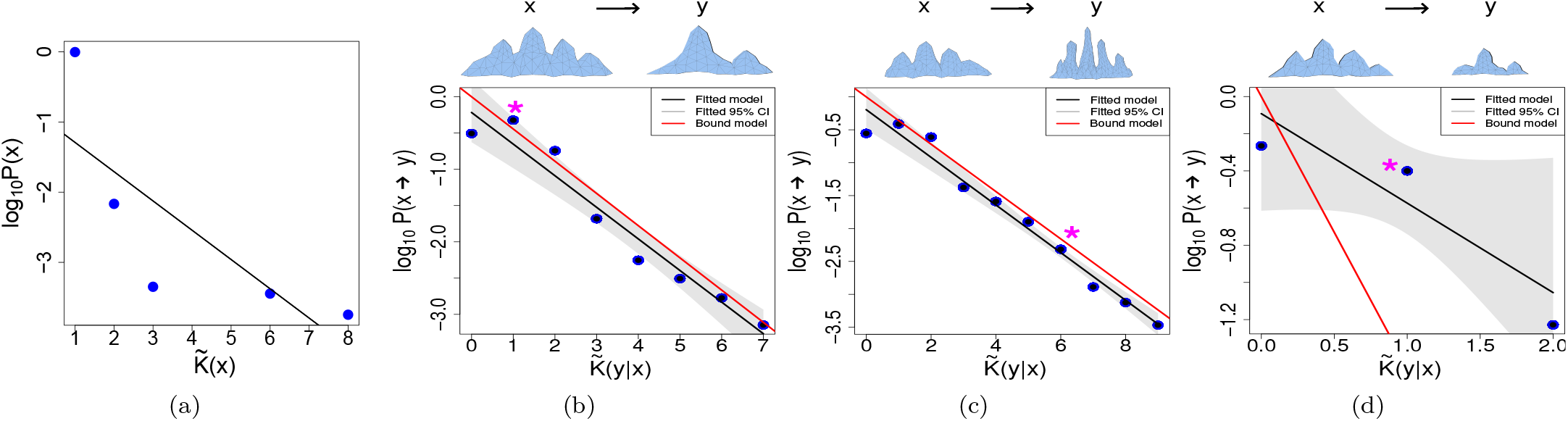
Tooth model with complexity as cusp number. (a) Simplicity bias found for the uniform random sampling of the tooth model where we measure complexity as the number of cusps. The black line shows the linear regression. On top of each of the following plots we can see the phenotype x on the left side and an example of a phenotype y, into which phenotype x has mutated. On the plots, the * indicates where the example y can be found. As previously explained, the following examples illustrate varying levels of success corresponding to different degrees of simplicity bias. (b) Example for the transition probabilities of a phenotype with complexity 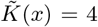. In this example *ρ* = − 0.20, such a negative correlation confirms Level (I). Level (II) is also achieved with an *R*^2^ = 0.95. Finally, Level (III) is also achieved, since the differences of the slopes calculated with the bootstrap method and the slope of the bound model are not significantly different from 0. (c) In this example, using a phenotype of complexity 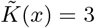, Level (I) is achieved with a *ρ* = − 1.0. Level (II) is also achieved, with an *R*^2^ = 0.97, Level (III) is also achieved. (d) In this example, using a phenotype of complexity 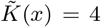, Level (I) is achieved with a *ρ* = − 1.00. Level (II) not achieved with an *R*^2^ = 0.79 however, Level (III) is not achieved, since the slopes of the fitted model and the bound model are significantly different.

The results for teeth using OPCR complexities are plotted in Figure 4. As can be seen in Figure 4(a), the tooth model with this new complexity measure likewise exhibits simplicity bias, as teeth with lower patch counts occur much more frequently in our exploration. In Figure 4(b-d) we generate three *in silico* teeth with complexity values of 86.25, 46.13, and 103.75, respectively, and introduce point mutations on single parameters within biologically realistic ranges. Mutants in Figure 4(b-d) exhibit Level I conditional simplicity bias: there is decay, and since Level II is also achieved in all cases, this decay is mostly linear. However, in 4(c-d) Level (III) is not achieved. In Table II we show that only in 34% of all the studied cases is Level III achieved, since the slopes are correctly predicted by the upper bound model. Notice that here, because the phenotype categories are not as distinctly separated, we divide the conditional complexity into 10 groups and select the point in each group with the highest transition probability to perform the bootstrap (the black points in 4(b-d)). Otherwise, the procedure to determine if Level (III) was achieved is the same as explained in Section III-C.

**FIG. 4.**
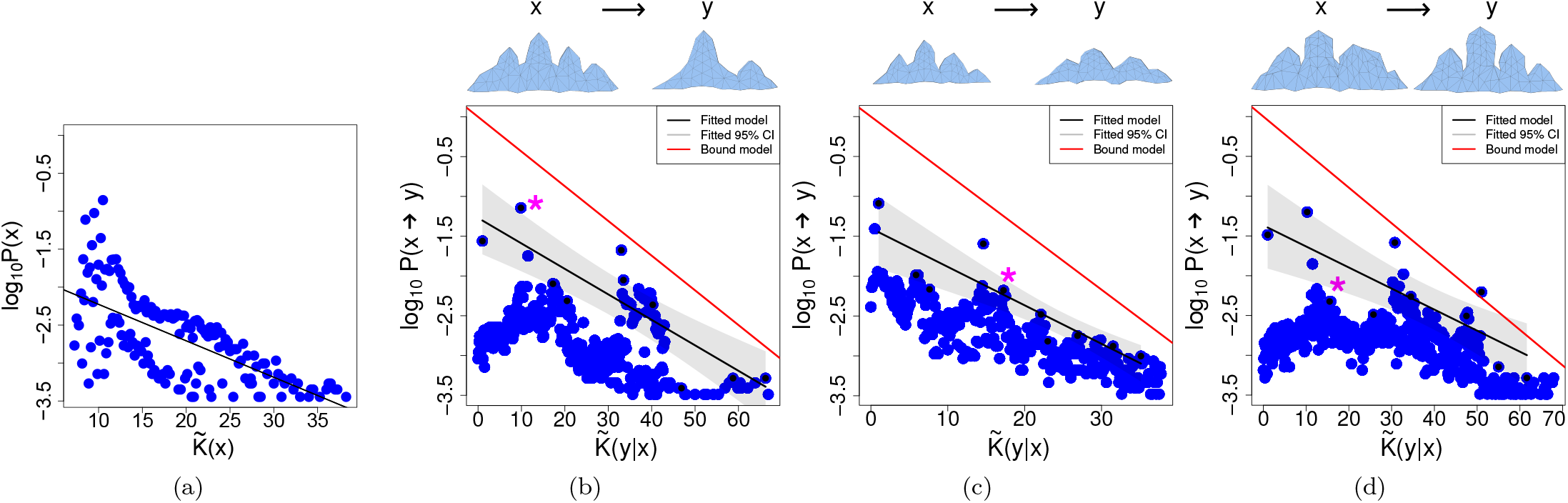
Tooth model with OPCR as complexity. (a) Simplicity bias found for uniform random sampling of teeth using OPCR as the measurement for complexity. The black line shows the linear regression using the maximal values for each category of complexity. On top of each of the following plots we can see the phenotype x on the left side and an example of a phenotype y, into which phenotype x has mutated to. On the plots, the ∗ indicates where the example y can be found. As previously explained, the following examples illustrate varying levels of success corresponding to different degrees of simplicity bias. (b) Example for the transition probabilities of a phenotype with complexity 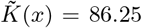. In this example *ρ* = − 0.45, this negative correlation confirms Level (I). Level (II) is also achieved, with a *R*^2^ = 0.74. Finally, Level (III) is also achieved, since the differences of the slopes calculated with the bootstrap method and the slope of the bound model are not significantly different from 0. (c) In this example, using a phenotype of complexity 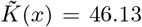, Level (I) is achieved with a *ρ* = − 0.80. Level (II) is also achieved, having a *R*^2^ = 0.78, however, Level (III) is not predicted, as slopes of the bound and fitted model are significantly different. (d) In this example, using a phenotype of complexity 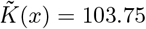, Level (I) is achieved with a *ρ* = − 0.30. Level (II) is achieved, with an *R*^2^ = 0.64, finally, Level (III) not achieved.

Taking advantage of a complex model that produces realistic shapes, we conclude that complexity of the mechanics of a generative system may not cause the simplicity bias to be weaker. This is corroborated by the fact that we see similar results irrespective of the complexity measure and even when applied to another species (Appendix C for the results using the tooth model adapted for sharks). Overall, the progressive rarity of complex tooth shapes in the tooth model does not come as a surprise. It is due to the fact that many developmental parameters need to be fine-tuned in order to achieve some level of phenotypic complexity and stability [71]. Additionally, there are always multiple ways that parameter changes can lead to failure in reproducing a phenotype, resulting in an unavoidable bias towards simpler shapes [78]. Notably, this theoretical argument has been corroborated experimentally [79]. As noted before, even though the map here is highly complex, it is not completely random. The tooth model follows some biomechanical rules and was conceived to be able to reproduce the natural variation found in seals [72], and in a more recent version, sharks [75]. This involved a choice of tunable mechanisms which was informed by knowledge about tooth development. Therefore, the interactions between the different components during development follow some non-random patterns, which are the result of an evolutionary process where some soft matter dynamics and biomechanical interactions are more likely than others [70]. This is quite different from the matrix multiplication map, where all interactions are completely random, leading to a medium complexity map (discussed above), so that the output may depend more of the map itself than of the input information.

Despite the generally monotonic decrease in the conditional probability of occurrence with increasing complexity, we have observed that this decrease becomes smaller towards the right side of the diagrams. This may be interpreted in terms of a relatively high robustness, meaning that more complex teeth are particularly likely to reproduce themselves following mutations. This may reflect disparities in the effects of different parameters on phenotypic changes, especially since the mutation step size was set to be gradual, allowing only negligible changes in values. Alternatively, the iso-morphological walk can be considered a proxy for an evolutionary process that is more likely to discover robust phenotypes within the morphospace. In addition, our observation may reflect another potentially general property of genotype-phenotype maps: different complex phenotypes tend to be clustered in islands within morphospaces, facilitating transitions between them [78]. This suggests that mutants with very complex phenotypes might either be extremely simple due to failed development, or, more often than expected, only slightly less complex, thus influencing the shape of the conditional probability function.

One possible caveat lies in the arbitrary end point of development, which excludes several *in silico* teeth whose complexity unfolds too slowly. Although this issue would arise with *any* choice of endpoints, it may partially explain the noticeable differences between the frequency of the simplest (1-cuspid) and all other complexity categories. Since development never decreases complexity, only the simplest category will remain unaffected by endpoint choices.

Interestingly, our results do not seem to be strongly affected by how complexity is discretized. Therefore, we suggest that the challenge of selecting the most appropriate complexity measure and data discretization method may not be the primary obstacles in quantifying complexity biases in biologically relevant complex traits.

### E. Polyominos

Polyominos are 2D square lattice tile shapes, formed of self-assembled individual square blocks [92, 93]. Each individual square block has labelled edges, with certain labels allowed to stick to certain other labels, and certain labels prohibited from sticking to certain other labels. In this genotype-phenotype map, the genotype specifies the rule-set determining which edge type can stick to which other edge type, and the overall multi-tile shape of the self-assembled polyomino defines the phenotype. For example, a single square block with no bonds is a (fairly trivial) phenotype; and a two-by-two square is an example of another phenotype. With a large number of tile-types and many tiles, a whole array of different phenotype shapes can be formed. Figure 5 shows some example 2D tiles shapes.

**FIG. 5.**
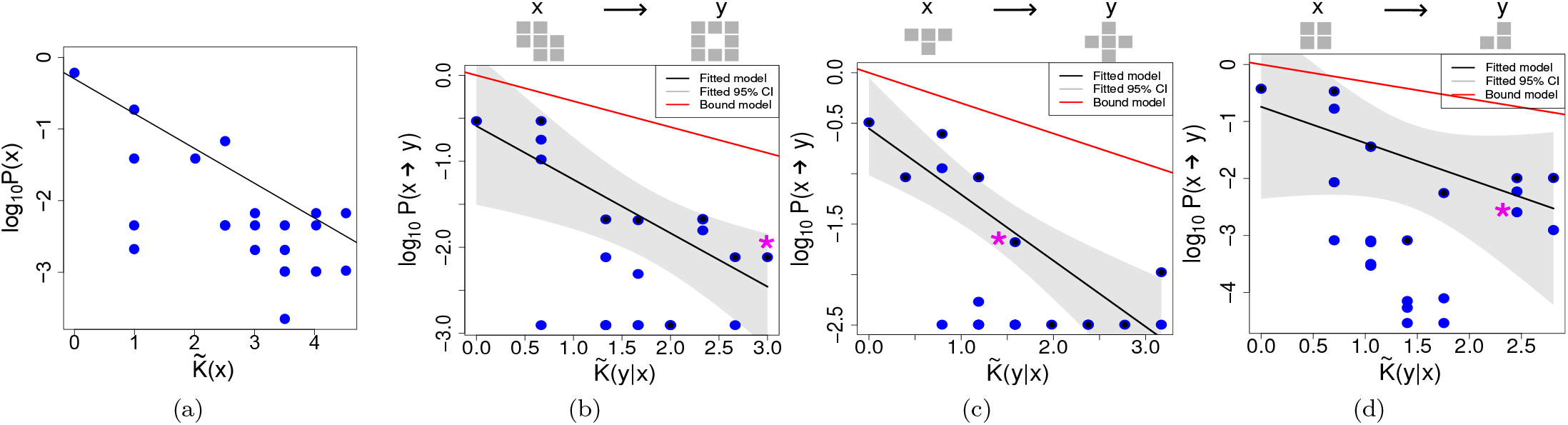
Self-assembling polyomino tiles (site-occupation). (a) Simplicity bias found for the uniform random sampling of polyominos. The black line shows the linear regression using the maximal values for each category of complexity. On top of each plot we can see the phenotype x on the left side and an example of a phenotype y, into which phenotype x has mutated to. On the plots, the * indicates where the example y can be found. As previously explained, the following examples illustrate varying levels of success corresponding to different degrees of simplicity bias. (b) Example for the transition probabilities of a phenotype with complexity 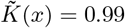. In this example *ρ* = − 0.26, such a negative correlation confirms Level (I). Level (II) is also achieved, with a *R*^2^ = 0.62. Finally, Level (III) is also achieved, since the differences of the slopes calculated with the bootstrap method and the slope of the bound model are not significantly different from 0. (c) In this example, using a phenotype of complexity 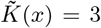, Level (I) is achieved with a *ρ* = − 0.58. Level (II) is also achieved, with a *R*^2^ = 0.75, however, Level (III) is not predicted, as slopes of the bound and fitted model are significantly different. (d) In this example, using a phenotype of complexity 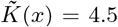, Level (I) is achieved with a *ρ* = − 0.29. However, Level (II) is not achieved, with *R*^2^ = 0.42, failing at this Level (II) also means that Level (III) is not achieved.

Despite the abstract nature of this genotype-phenotype map model, polyominos have been used to model biological self-assembly, for example in terms of protein quaternary shapes [93], including to successfully explain certain aspects of protein evolution [30, 94]. The polyomino model we use here has a genotype which is a binary string specifying which tiles faces can stick to which other tile faces. The data set comes from ref. [94] (specifically, the *S*_2,8_ data), and there are 22 different phenotype shapes.

The polyomino map has been examined in terms of simplicity bias in the sense of Eq. (2), and it was shown that clear simplicity bias is observed [30]. However, the complexity measure used in that earlier study was designed with polyominos in mind, rather than being a completely generic map-agnostic complexity measure for 2D tile shapes. Naturally, a complexity measure designed with the specific map in mind will likely produce more accurate probability estimates or bounds than a generic complexity measure. However, if the goal is to try to create an information complexity theory that applies at least somewhat to a whole range of maps without having to know the details of the mapping process, then using a map-specific complexity measure is not ideal.

With this goal in mind, here we utilise a generic complexity measure applicable to polyominos, which we can call *site-occupation complexity*. It can be used to study both simplicity bias and also conditional simplicity bias in polyominos. The site-occupation method is as follows: Because polyominos consist of square blocks on a grid, we can represent their shapes by placing a ‘1’ in a grid site if there is a block, and a ‘0’ otherwise. In this manner, any polyomino can be represented by grid site occupation. The resulting 2D binary grid can then be represented as a 1D string by concatenating rows of the grid, and then the complexity value of each polyomino can be computed using *C*_*LZ*_ and Eq. (5). For example, a hollow four-by-four square would be read as: 1111 1001 1001 1111. Similarly, conditional complexity can be calculated by concatenating 1D strings, following Eq. (4), in order to make the transition probability predictions, *P* (*x → y*).

Figure 5(a) shows simplicity bias in the polyomino model. Turning to panels Figure 5(b), (c), and (d) we see that a roughly linear upper bound decay appears, but the slope is not accurate. Hence we have achieved Level (II) for this map. See also Table II. In Appendix D we show how a different measure, based on the complexity of the perimeter of the polyomino shape, also yields comparable simplicity bias and transitions plots.

### F. HP proteins

A protein sequence genotype folding to the 3D protein tertiary structure phenotype is a well-studied genotype-phenotype map. For natural proteins this map has been hard to study due to the (until recently) unsolved problem of directly predicting the structure from the sequence, and also due to the great variation in natural protein sizes, architectures and functions. Due to these issues and others, many studies of simplified protein maps have been undertaken, with the goal of uncovering the key aspects of this sequence-structure map. These simplified models still have relevance despite the recent success of machine learning algorithms in predicting structures, because while such algorithms can yield accurate predictions, they do not help to the same degree in a theoretical understanding of maps themselves, their properties, and basic physics.

A popular model in computational studies of genotype-phenotype maps is the HP protein model [95, 96]. In this model, the process of protein folding is simplified: Firstly, in the model there are only two types of amino acid, labelled as either hydrophobic (H), or polar (P), instead of the full suite of 20 amino acids present in nature. Secondly, the protein structures are confined to a 2D lattice so that there are only finitely many possible 2D structures for a given length HP protein chain (note that 3D lattice models also exist [97]). Sequences ‘fold’ to their minimum energy structures, where energy values come from counting nearby energetically favourable interactions in the chain. In this model, if a given sequence has two or more different shapes with the same energy, then the sequence is said to have no structure and is discarded. This requirement is supposed to model the natural phenomenon that when a protein sequence does not have a well-defined and stable tertiary structure, it may not be viable in an organism. Some example HP proteins are depicted in Figure 6.

**FIG. 6.**
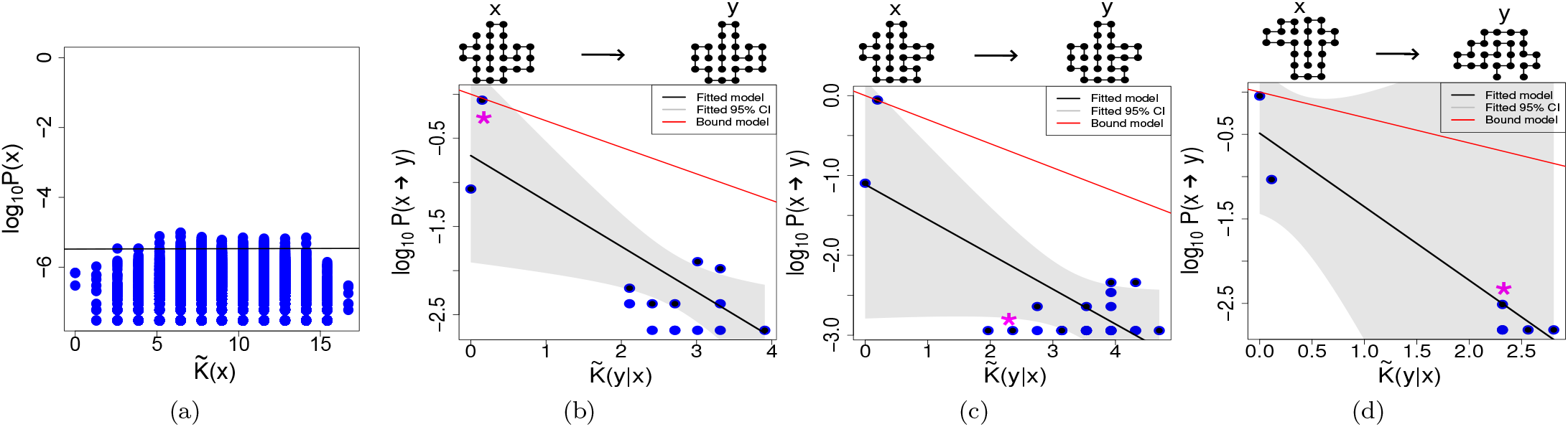
The HP protein map. (a) No simplicity bias found for the uniform random sampling in the HP protein map. The black line shows the linear regression using the highest probability values for each unique complexity value. On top of each plot we can see the phenotype x on the left side and an example of a phenotype y, into which phenotype x has mutated to. On the plots, the * indicates where the example y can be found. As previously explained, the following examples illustrate varying levels of success corresponding to different degrees of simplicity bias. (b) Example for the transition probabilities of a phenotype with complexity 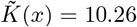. In this example *ρ* = − 0.45, this negative correlation is suggestive of achieving Level (I), but a visual inspection of the data highlights that in fact there is little evidence of a trend. The metrics suggest Level (II) and Level (III) are also achieved, with a *R*^2^ = 0.73 but again visual inspection of the data substantially reduces our confidence in these conclusions. (c) In this example, using a phenotype of complexity 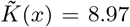, Level (I) is apparently achieved with a *ρ* = − 0.10, but the same comments as for a (b) apply. Level (II) is also apparently achieved, having a *R*^2^ = 0.53, and even Level (III) is achieved according to the metrics, but the visual inspection of the data implies that the evidence of conditional simplicity bias is weak. (d) In this example, using a phenotype of complexity 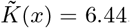, we have *ρ* = − 0.84 and a *R*^2^ = 0.93, while Level (III) is not achieved. Again, the paucity of data and lack of intermediary complexity and probability points makes it hard to draw conclusions.

It is well known that the number of genotype sequences is much larger than the number of possible phenotype HP protein patterns [95]. Additionally, the resulting phenotype is a discrete chain on a lattice, and this chain can be described (for example) as a chain of directions on a lattice: up (U), down (D), right (R), and left (L). In this manner, a 2D HP lattice protein can be represented as a string of characters (U, D, R, L), and a complexity value for the structure can be computed using *C*_*LZ*_ and Eq. (5). Conditional complexity estimates can be obtained also in this manner, via strings of characters. Naturally, there are other ways that the lattice protein could be described also, but this intuitive and precise method will suffice us. In this genotype-phenotype map, the connection between the inputs and outputs is direct, similarly to the RNA and protein secondary structure models also studied earlier [33], but this model has not been studied for simplicity bias nor for conditional simplicity bias before, and so is examined here.

Because earlier studies of HP protein maps have reported phenotype bias, and at least one example where the structure with the highest probability displays overall symmetry [95], we might expect to see simplicity bias in the HP protein model. To test for simplicity bias for the HP protein model, we will use data from ref. [96] which has sequence length of *L* = 25. Note that in this model, all self-avoiding walks on the 2D lattice are considered potential phenotypes, and not only maximally compact structures (which are sometimes restricted to). Despite expectations, in Figure 6(a) we see that there is no simplicity bias observed. It is not completely clear why this is, but we can propose some possible reasons: Firstly, it may be that the *C*_*LZ*_ complexity measure we use is not able to detect the relevant patterns in HP proteins. Secondly, it could be that the structures we use are too small to show clear simplicity bias (it was shown earlier than simplicity bias only emerges in RNA secondary structure for longer sequences [32]). Thirdly, it is apparent that there is relatively little *bias* in this map, because the probabilities vary only over one or two orders of magnitude, despite the broad range in complexity values. Clearly, without strong bias (i.e., strongly non-uniform probabilities), there cannot be pronounced simplicity bias (see [56] for a discussion and example of this). See below for more on the strength of bias. Relatedly, there do not appear to be any HP protein structures that have very high probability.

Now turning to consider conditional simplicity bias, we employ the same data set and plot the conditional complexity graphs in Figure 6(b), (c), and (d): the data do not show clear evidence of conditional simplicity bias. In each panel, there are two clusters, one at high probability and low conditional complexity, and the other at high complexity and low probability. However, between these two clusters there is an absence of intermediary complexities and probabilities. Hence these data are inconclusive, and do not provide strong evidence of conditional simplicity bias. It is worth highlighting that unlike in panel (a), panels (b), (c), and (d) show large variations in probability (∼ 3 orders of magnitude). There is stronger bias here even while there is not much bias in panel (a). We can conclude that there is some very modest evidence of observing Level (I), but no clear conclusions can be made regarding Level (II) and Level (III).

To briefly explore another HP model, we used data for the *L* = 36 HP protein (code from [98]), which studies the compact version of the HP model, where only maximally compact phenotypes are accepted as potential phenotypes. This compact model may potentially have different map properties, and hence it is interesting to study this model too. However, even using this model, the resulting plots are very similar, with weak or inconclusive evidence of simplicity bias (see Appendix Figure D.3).

## V. DISCUSSION

### A. Summary of findings

We have investigated an approach to predict, or at least bound, the probabilities of phenotype transitions upon random genetic mutations, using arguments inspired by algorithmic information theory, and especially the phenomenon of conditional simplicity bias (Eq. (3)). Earlier [33], it was shown that the transition probabilities in computational simulations of RNA and protein secondary structure genotype-phenotype maps could be upper-bounded by estimating the complexity of the starting and resulting phenotypes. The ability to make such predictions was noteworthy because it suggested that map-agnostic bounds, just relying on information complexity arguments, could provide non-trivial predictions of transition probabilities, which may be useful in cases where the details of the underlying genotype-phenotype map are not known. More broadly, the ability to make such predictions supported the exploration of information complexity arguments for developing mathematical laws in biology. Further, such predictions support understanding the nature of biases in the introduction of phenotypic variation, which may be relevant to evolutionary dynamics.

In the present study, we have extended this research direction by applying the conditional simplicity bias bound to several other genotype-phenotype maps, and in particular more ‘challenging’ maps were chosen which in one way or another tested the limits of the applicability of the conditional simplicity bias bound. These included a differential equation model of a circadian rhythm, a matrix map, a detailed tooth development model (with two types of complexity estimate), a polyomino self-assembled protein complex map, and an HP lattice protein map. The fact that the model of teeth development — which is highly intricate and biologically realistic — shows conditional simplicity bias (and simplicity bias), is noteworthy because it suggests that the bound may be applicable at higher or other levels of biological organisation.

Overall the numerical experiments show that (i) conditional simplicity bias appears in most of these maps, and (ii) some degree of transition probability predictability can be achieved, varying between maps. In nearly all cases, Level (I) conditional simplicity bias was achieved, meaning some general inverse relation between probability and complexity. In several cases, Level (II) was achieved in which the upper bound on log *P* (*x* → *y*) was found to be roughly linear. Not many cases achieved Level (III), in which the slope was also correctly identified.

### B. Possible reasons for reduced prediction accuracy

There are several possible reasons which might explain why not all levels were achieved in all maps. Firstly, the complexity measure may not be suitable for all maps: It may be that the *C*_*LZ*_ method cannot handle all types of patterns that occur, and it is already known that pseudo-random patterns cannot be handled by this method [99, 100], for example. It is possible that, say, in the HP proteins the relevant patterns cannot be detected. Exploring more powerful pattern detection and complexity methods may help with this. Secondly, it has already been noted [32] that simplicity bias emerges only for large enough systems, when finite-size effects do not dominate the patterns and trends. Hence it is possible that for larger system sizes, more clear and convincing (conditional) simplicity bias may be observed. On the other hand, the computational requirements are taxing in this case. Thirdly, the simplicity bias bounds are most applicable when the underlying maps exhibit strong bias, and so if the maps do not exhibit strong bias then this may affect the upper bound predictions. Fourthly, as discussed above, the Level (III) predictions depend on having good estimates of the number of accessible phenotypes and the maximum and minium complexity values. Because these values are sometimes hard to estimate, this can reduce the accuracy of the slope predictions. Fifthly, the slope of decay can be affected if the phenotypes/output are partially randomised [57]. It could be that in some maps, the slope is affected by similar factors. Finally, the underlying theory for the slope predictions may need to be developed further, for example, in a way which takes other phenotype factors into consideration, beyond just the conditional complexities (e.g., the complexity or probability of the starting phenotype *x*).

### C. Limitations of predictions and simplifications assumed

A weakness of our predictions is that they only constitute an upper bound on the probabilities, with many phenotype patterns falling far below their respective upper bounds. These phenotypes have low probability values, while at the same time low (conditional) complexity values. Following the hypothesis from refs. [52, 58], these low-complexity low-probability outputs are presumably patterns which the genotype-phenotype map find ‘hard’ to make even though they are not intrinsically very complex. It may be possible to extend approaches developed in [52, 58] that also take into account the complexity of the genotypes to help explain which types of patterns occur far from the bound, and to find improved estimates for their probabilities. Having described this weakness in predictions, from a different perspective the problem is not as severe as it might seem: It is known that randomly sampled genotypes are likely to generate phenotypes which are close to the bound [52]. In other words, even if many of the phenotypes are far below the bound, most of the mutations map to phenotypes which are close to the bound. A related weakness is that we can rarely predict the value of the slope of the upper bound, that it, Level (III) was rarely achieved. Even with this weakness, we can still predict other properties of interest, like for example which of two phenotypes is more likely [33].

In this study we assume uniformly random mutations which ignores mutational biases [20] at the genome level. While this is a simplification of reality, it is unlikely that incorporating these biases would drastically alter the global patterns of transition probabilities we demonstrated here, see e.g. [28] for an example. Nevertheless, there may be interesting directions to study where mutational biases and phenotypic biases interact. Similarly, the effect of compositional bias in genomes such as CG bias, may affect phenotypic biases.

As alluded to above, other methods for predicting phenotype transition probabilities, in particular those which invoke biophysical details of the map, or other details of the relevant genotype-phenotype map, will no doubt yield more accurate predictions for transition probabilities (e.g., [98]). Nonetheless, these other methods have a different aim and different list of requirements and assumptions, and hence cannot be meaningfully compared to the predictions done here.

### D. The meaning of complexity

The word “complexity” can take on many meanings in the scientific literature [78, 101–105]. While we use the concept of complexity, and are applying results inspired by Kolmogorov complexity to biology, we are not in anyway trying to argue that one or other approach to measuring complexity in biology is more or less valid. The aim of this work is not primarily to measure complexity, but rather to use a body of theory that provides methods for predicting probabilities, which uses a particular form of complexity to achieve this goal.

While strictly Kolmogorov complexity is uncomputable, it can be estimated using methods such as data compression, or other computable measures of descriptional complexity [49]. In practice we have mainly employed the *C*_*LZ*_ complexity measure which is related to lossless compression techniques, and which is a standard and theoretically motivated approximation to the true uncomputable quantity. We have also employed other descriptional complexity measures here, i.e. cusp-count and OPCR. The former is a biologically intuitive but quite coarse-grained measure of the descriptional complexity for teeth shape, while the latter is a more fine-grained descriptional measure the variability of tooth’s surface. Previous work has shown that well-motivated approximations to descriptional complexity can work well in capturing the biases predicted by the AIT derived bounds [30, 32, 53–55].

### E. Implications for robustness, evolvability, and fitness landscapes

A phenotype is said to be robust to mutations if the probability of changing to another phenotype upon random genetic mutations is relatively low [65]. According to the derived bound on *P* (*x* → *y*), point mutations are a priori more likely to change a shape *x* to a shape *y* which is (i) very similar to *x*, or (ii) a very simple/trivial pattern (while in this case not necessarily similar to *x*). Because the most similar shape to *x* is *x* itself, the authors of ref. [33] pointed out that this conditional bound naturally gives rise to high genetic robustness as a null model for genotype-phenotype maps (under some conditions), while the origin of the high robustness in genotype-phenotype maps had previously been seen as something of a mystery [106, 107]. Mohanty et al. [108] recently mathematically derived upper limits on the possible robustness of phenotypes, and high robustness was linked to phenotypes whose genotypes mainly consist of sections of constrained and unconstrained regions (see also [106]). While such architectures will lead to high robustness, it is interesting that the conditional complexity bound does not explicitly invoke this, and so may predict robustness as a null model when such architecture is not present or known to be present.

Evolvability is a concept which refers to the ability of populations to heritably change and adapt in reaction to changes in selection pressures [65, 109, 110]. In order to make such adaptations, a given phenotype must have a variety of phenotypes which are mutationally accessible. In the nomenclature of this paper, it means that *P* (*x* → *y*) must be positive for many different *y*. There is some tension between robustness and evolvability [109], because one appears on the face of it to preclude the other although some theories of how to reconcile these concepts have been proposed [111, 112]. Intriguingly, the conditional complexity upper bound may point to another resolution to this tension by (1) giving a high (null-model) transition probability to *y* = *x* (high robustness), and high probability to *y* which are similar to *x* (a kind of generalised robustness), and yet at the same time (2) allowing for many different *y* to have non-zero transition probabilities thereby aiding evolvability.

There may also be extreme cases of zero-probability transitions occurring such that *P* (*x* → *y*) = 0, indicating no direct mutational pathways between two phenotypes. Such connections are crucial for navigating fitness landscapes, because they determine the accessibility of evolutionary pathways [113]. The navigability of a fitness landscape can be analysed in terms of a directed phenotype network [114]: When most phenotypes are connected, navigability is high due to potential high-dimensional bypasses. In contrast, if most phenotypes lack connections, the fitness landscape becomes rugged, making it difficult for evolving populations to locate fitness peaks.

Additionally, it is reasonable to assume that fitness differences may also be linked to conditional complexity. Phenotypes with lower conditional complexity relative to a high-fitness phenotype are likely to exhibit higher fitness, while the opposite may also hold true. This effect would suggest that larger transition probabilities are typically towards phenotypes with more similar fitness, generating interesting correlations in fitness landscapes, and potentially causing the relevant fitness landscapes to be smooth. All this suggests that finding genotype-phenotype map agnostic information theory arguments to predict the topology of a genotype-phenotype-fitness map may be a fruitful future research programme. We leave for later works the task of exploring in depth of how conditional simplicity bias relates to robustness, evolvability, navigability, and fitness.

### F. Evolutionary dynamics and adaptive landscapes

Here we consider how the simplicity biases studied in this work might interact with other evolutionary processes. Since we have been strictly concerned with the relationship between genotypes and phenotypes without evolutionary change, our models can be useful by representing general null-models to contrast variation reflecting developmental biases and the variation in population as a result of evolution. Unlike *in silico* approaches, real natural populations are the outcome of different kinds of selection and may therefore show biases different from the null-model predictions. As a concrete example, we discuss the tooth model where ample data from extant populations are available. First, developmental studies have demonstrated that both experimentally interfering with odontogenesis and mutagenesis are more likely to simplify tooth shapes than to lead to more complex teeth, providing evidence in favour of the proposed simplicity bias [79, 115, 116]. Conversely, it has been suggested that throughout evolution, dental complexity has increased in most mammalian clades [81, 117, 118], with some notable exceptions [119]. This shows that developmental and evolutionary biases are not necessarily aligned, with functional and ecological pressures often dominating long-term phenotypic changes. Yet, while non-alignment of biases is widespread, phenotypic change is likely to be accelerated if aligned with the direction of developmental biases, or possibly slowed down, in the contrary case. For instance, there is evidence that the evolution of complex molar types may take unusually long [120] (although comparisons between rates of evolution have to be considered with caution as they as they depend on various factors such as size and proliferation rate). Cases of dental simplification have been suggested to result from relaxation of functional or ecological constraints, which is precisely what our hypothesis would predict [119, 121].

A common framework to understand and visualize the evolutionary dynamics of variation that is used in population genetics is the adaptive landscape metaphor [122, 123]. Therein, fitness is mapped onto genotypic or phenotypic traits. Assuming a situation where high complexity is associated with higher fitness, we would find that fitness peaks become increasingly (with altitude) rugged, steep and rare, while smooth valleys abound. Any population starting from a point on the slopes would be more likely to fall down into such a valley than climbing uphill, and increasingly so along their evolution. Strong selection may, thus, be needed to sustain their trajectory against the theoretical tendency towards relapsing into lower territories. In mammals, dietary function and occlusion may be such selection pressures. By linking complexity and landscape topology, our study is well-nested within long-standing theory of the evolution of populations which may allow for quantitative study designs. Thus, by systematically testing whether general developmental biases exist, we aim to identify generative null-models which can be meaningfully compared to biological data which are the intertwined outcome of developmental and evolutionary processes.

### G. Biases

Interest in the topic of biases in the introduction of variation has grown in recent years [4, 11, 20]. This recognition contrasts with a common (even if tacit) assumption that variation is either roughly uniform, or that biases do not have much impact on evolution [11]. It also contrasts with the view that biases merely limit certain possibilities [9], as opposed to positively affecting direction of evolution [124, 125]. Within this framework, we see that simplicity bias and conditional simplicity bias provide further sources of what can be quite a pronounced bias in the introduction of variation.

Related to this, Salazar-Ciudad [126] raises the interesting objection that the term “bias” is improper: there is no bias, but simply the action of development. The argument being that it is not correct to first imagine possible phenotypes in a morphospace, and then react with surprise when these imagined possibilities do not manifest, or only a small fraction of these possibilities appear. While we agree that there is a valid point raised here, especially in the context of development, we maintain the appropriateness of the word “bias” in our context. This is due to the fact that we use bias to mean that assuming a uniform distribution of random mutations, a non-uniform (biased) distribution over phenotypes will result, and that this bias has certain predictable properties, in particularly a prefernce for simplicity. This is the sense in which we mean simplicity “bias”.

It is interesting to open the discussion regarding whether or not bias itself is an evolved property [11, 125]. Some questions are: Have biases changed over time? Can biases in genotype-phenotype maps be tuned via evolution? It is conceivable that some biases may have adapted to the needs of the organism; that is, it may be that the genotype-phenotype map in some cases adapts itself via evolution so that the phenotypes which are ‘favored’ by bias are the ones which are most needed to adapt to the environment. In many biological contexts this type of adaptive argument is plausible, but as argued by Ghaddar and Dingle [31], there are other types of bias which result from basic physical, chemical, or information constraints. In these cases, it is difficult to see how selection could alter these fundamental properties, which would be needed to alter the biases. This observation applies here also to the case of biases arising from conditional simplicity bias. Having said that, one area in which there is still potentially room for adaptation to tune the bias is in the context of low-complexity, low-probabilities phenotypes [52, 58]. The information constraints apply to the upper bound probability, but not directly to how far a given phenotype’s probability is below the bound. Hence there could conceivably be some tuning of the bias in relation to the distance from the upper bound.

Concluding the discussion of biases, it is interesting that unlike other developmental biases, the biases arising from (conditional) simplicity bias do not depend on evolutionary history, making it easier to predict the direction and type of bias. This follows from the fact that (conditional) simplicity bias derives from intrinsic information arguments, and the relative simplicity or complexity of phenotypes, and these quantities can be estimated a priori. This predictability contrasts with other developmental bias cases like the famous example of patterns of digit reduction in amphibians [127]. These patterns of digit reduction observed were (presumably) contingent, and not predictable a priori, but instead could only be uncovered via direct observation.

### H. Future directions

Looking to future work, three directions stand out. Firstly, there is the question of how the link between conditional complexity, transition probabilities, and fitness differences affects the topology of fitness landscapes. Secondly, it would be interesting to study transition probabilities in other biological models such as plant branching patterns (via L-systems) [128], biomorphs [17], and leaf shapes [129]. The third direction is more mathematical: for the basic simplicity bias described by Eq. (2) we can predict the slope quite well [32], but in the conditional simplicity bias plots which employ Eq. (3), our ability to predict the slope is not as good (as we have seen here in the current study). Can we explain the origin of this discrepancy, and perhaps find better predictors? Finally, while the current study has focused on genotype-phenotype maps, which is a biological context, the conditional simplicity bias bound in Eq. (3) should be a generic property of input-output maps, and be applicable beyond biology, such as in genetic programming [130–132]. Hence there are potentially many applications and extensions for this line of research.

## Acknowledgments

This project has been partially supported by Gulf University for Science and Technology and the CAMB research center under project code: ISG Case 9 and ISG Case 44. We thank Iain Johnston and Sam Greenbury for providing polyomino data.

## Data availability

The code and data used in this work are available at GitHub: https://github.com/hagolani/Bounding-phenotype-morphology-transition-probabilities-via-conditional-complexities

## Appendix A: Applications of Kolmogorov complexity

The application of AIT to real-world science problems suffers from several problems, including that (a) Kolmogorov complexity is uncomputable, (b) the results are framed and proved in the context of universal Turing machine (UTMs) while many real world maps are not Turing complete, and (c) results are valid up to *O*(1) terms and therefore, strictly, only accurate in the asymptotic limit of large complexities. Given these, it is surprising that AIT and algorithmic probability can be successfully applied at all. However, several lines of reasoning can help us understand why they are, in fact, applicable, at least approximately.

Firstly in response to (a), although technically uncomputable, Kolmogorov complexity is fundamentally merely a measure of the size in bits of the compressed version of a data object. Hence, Vitanyi [48, 133] points out that because naturally generated data is unlikely to contain pseudo-random complexities like *π*, the true complexity is unlikely to be much shorter than that achievable by every-day compressors. For more on this discussion, see ref. [133] and ref. [134] for work on short program estimates via short lists of candidates with short programs. Secondly, in response to (b) it is worth noting that the simplicity bias bound is specifically relevant in the computable (i.e., non-UTM) setting. Moreover building on pioneering work with small Turing machines [62, 63], Zenil et al. [135] have numerically studied algorithmic probability at different levels of the Chomsky hierarchy and found that it persists. Indeed, they also found close agreement between complexity estimates obtained from different levels, for the short binary strings which they studied. From the theoretical side, Calude et al. [136] developed an AIT for finite state machines (the lowest on the Chomsky hierarchy), deriving analogous results, suggesting that many fundamental ideas and results from AIT need not only apply to UTMs. See also somewhat similar in ref. [137], and also the success of the Minimum Description Length (MDL) approach to statistics [138] which is a kind of computable version of AIT. Given that UTMs (at the top of the Chomsky hierarchy) and finite state machines (at the bottom of the Chomsky hierarchy) share many similar mathematical results related to AIT, it is not unreasonable to assume that the results hold for other systems (such as some biological systems) which sit between the two extremes of the hierarchy. Thirdly, we argued earlier [32] that it is common in physics that mathematical formulae apply quite accurately well outside the (e.g., asymptotic) regions for which they have been proven. Although it is not possible to completely remove *O*(1) terms in AIT [139], it is still an interesting question in theoretical computer science why asymptotic analysis (such as ignoring *O*(1) terms) is valid and works so well in practical applications [140].

Further reasons to understand why the AIT coding theorem should work in real-world applications can be found in information theory research developed largely independently of algorithmic probability. The fundamental connection between probability and data compression has also been studied by Cover [141], Langdon [142], and Rissansen [143]. Since then, different communities — e.g. information theory [144, 145], optimal gambling strategies [146], and password guessing [147] — have studied and exploited the probability-compression connection without explicitly invoking Kolmogorov complexity or UTMs.

In a review, Merhav and Feder [148] surveyed results in the area known as *universal prediction* and explicitly point to 2^−*LZ*(*x*)^ as an effective universal probability assignment for prediction based on the results of ref. [149] and others, where *LZ*(*x*) is the 1978 Lempel-Ziv compression complexity measure, essentially the same as we use here. Additionally, Merhav and Cohen [147] use a conditional coding theorem predictor 2^−*LZ*(*y*|*x*)^ which is again very closely analogous to the AIT conditional coding theorem relation, 2^−*K*(*y*|*x*)^ [64]. These results, together with the arguments given above in response to (a)—(c) all support and motivate the application of AIT coding theorem work in science and engineering, and also extending fundamental research related to the coding theorem and algorithmic probability [150].

The logic of this current study, and previous simplicity bias work [32, 33], is that we use AIT arguments to derive mathematical relations which can be proven to hold only in asymptotic limits, or under other conditions, but then apply these relations in other settings such as biology, and then simply test empirically whether or not the relations hold. Hence we have called this type of work “AIT-inspired arguments”, because it is perhaps not strictly AIT, which is a precise and abstract mathematical field.

## Appendix B: Map complexity

The complexity of the map is an important consideration in simplicity bias studies. It has been suggested [32] suggested that the map should be simple, or *O*(1) complexity. What the condition is imposing is that the complexity of the output patterns should be due to the complexity in the input, rather than the complexity (hidden) in the mapping procedure.

As an extreme example, if there were only two possible inputs, 0 and 1, and 0 mapped to the Shakespeare’s *Romeo and Juliet* while 1 mapped to *Hamlet*, then the complexity of the outputs (Shakespeare’s plays) would be due to the fact that the texts must be already programmed into the map. In this case, the inputs are very simple and in no way account for the complexity of the outputs. On the other hand, if a competent programmer writes many lines of code to produce a complex output via only a basic computer with few in-built functions, then the complexity of the output would be due to the complexity of the input program, and not merely the map itself.

It follows that if an arbitrarily complex map is permitted, then the map could be chosen so that complex outputs have high probabilities and simple outputs have low probabilities, or the assignments of inputs to outputs could be chosen in a way that there is no connection at all between complexity and probability.

## Appendix C: Tooth developmental model adapted for shark teeth

Here, instead of using the tooth model as in [72], we used a modified version adapted to shark tooth development [75]. Similarly to the original tooth model, in order to study the shark tooth model, we did some modifications to the strategy that has been followed for the other models presented here, mainly to account for the continuous parameters inputs, however this strategy differs from the one used in the tooth model, as we describe in the following paragraphs. We first identified one tooth for each of the following number of cusps *n*_*c*_ = 1, 3, 5, 7, 9, 11. Teeth with an even number of cusps were excluded since their appearance was rare due to the symmetric default development [151]. These initial teeth were taken from random morphospace explorations performed in [75]. In order to be able to use several different phenotypes with diverse parameter values, we then performed 100 iso-morphological walks [105], for each of these initial teeth. In each of the iso-morphological walks, teeth underwent 100 sequential steps of mutation, under the condition that their initial number of cusps remained unchanged in any tooth along the walk. Mutations are introduced by modifications of the model parameters. Specifically, for each mutant, we changed two randomly chosen parameters p to values ranging between the value of the respective parameter in the “parental” tooth *p*_0_ and either the minimal or the maximal allowed parameter value pm (as in [75]) in the following manner: *p* = *p*_0_ +(*α*_2_· (*p*_*m*_ −*p*_0_)) ; with *α* being a homogeneously distributed random number between 0 and 1. This procedure differed slightly from most other genotype-phenotype maps explorations in this study which typically introduce a single point-mutation at a time. However, we found that introducing two mutations was the most incremental way that allowed for an exploration of changes in dynamic parameter interactions and, thus, for an efficient detection rate of diverse tooth phenotypes. This also aligns more closely with natural evolution, where a small number of mutations often affects multiple developmental mechanisms. By performing these iso-morphological walks, we obtain 100 teeth per cusp number category *n*_*c*_. Finally, in order to measure the conditional complexity, we explored the two point-mutation neighborhood of each tooth in each category. Since we were using continuous values in this model, we could not explore every single possible mutation. Instead we explored 200 mutants per “parental” tooth. In order to account for these differences, when analysing the results for this exploration, we calculate the average frequency per complexity and conditional complexity category, since we are analysing the isomorphological space of cusps numbers rather than of specific phenotypes, since here we are defining the phenotype and complexity in the same way. In Figure C.1(a) we show that despite the added complexity the tooth model adds to the genotype-phenotype map, the probability of finding simple teeth is consistently larger than of finding more complex teeth, which is in line with other models and was to be expected. In fact, the decrease of the logarithm of frequency with increasing phenotypic complexity follows a near-linear curve, with the notable exception of mono-cuspid teeth, i.e. the lowest possible complexity. It could be argued that the linear probability decay may reflect that there is no qualitative difference between the mechanisms underlying an e.g. 3-to-5 and 5-to-7 cusp change. The high frequency of very simple shapes (1 cusp) might reflect that this minimum complexity exists for free, i.e.without the activation of specific mechanisms, and can be equally accessed from any part of the morphospace. As shown in C.1(b-d), the conditional simplicity bias appears to hold for the shark tooth model, with transition probabilities generally kept below the fitted and substantially below the bound model. Overall, the upper bound model appears to be a good predictor for the maximum transition probabilities. However, the monotony by which transition probabilities decrease appears weaker for more complex parental teeth. Level (I) and (II) are achieved in all the studied cases, which mean that we always see an linear fitted upper bound decaying with increasing conditional complexity. Even Level (III) is achieved in 73% of the cases.

**FIG. C.1.**
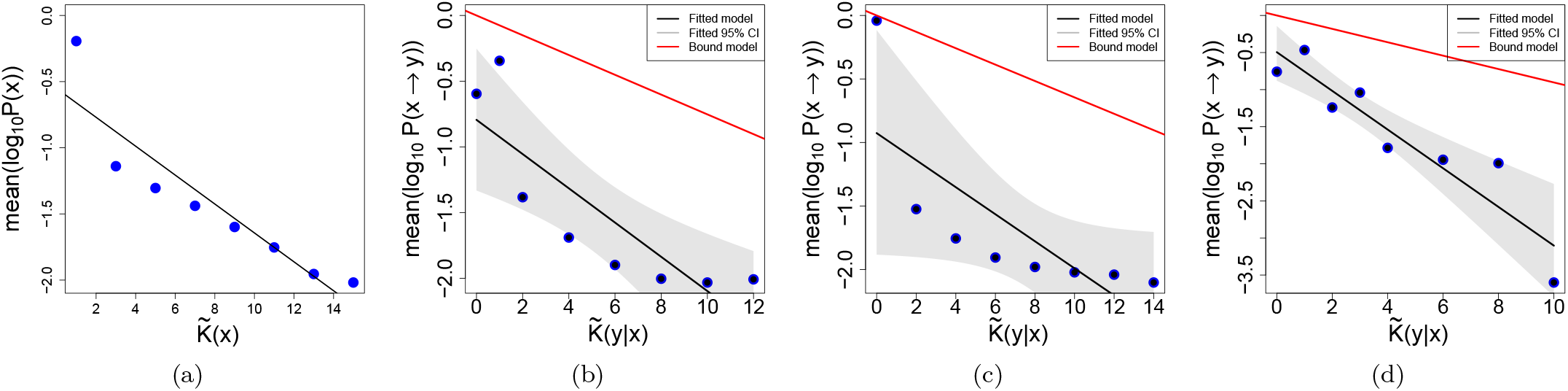
Shark tooth morphology model.(a) Simplicity bias found for the uniform random sampling of the tooth model where we measure complexity as the number of cusps. As specified in this section, this plot shows the average mean probabilities found for each of the complexity categories (number of cusps). (b) Example for the transition probabilities of a phenotype with complexity 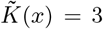. In this example *ρ* = − 0.95, such a negative correlation confirms Level (I). Level (II) is also achieved, since for the linear model built with this data, p-value=0.0077. Finally, Level (III) is also achieved, since the differences of the slopes calculated with the bootstrap method and the slope of the bound model are not significantly different from 0. (c) In this example, using a phenotype of complexity 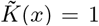, Level (I) is achieved with a *ρ* = − 1.0. Level (II) is also achieved, having the linear model a p-value=0.0281, Level (III) is also achieved. (d) In this example, using a phenotype of complexity 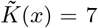, Level (I) is achieved with a *ρ* = − 0.95. However, Level (II) not achieved, since the linear models p-value=0.0009, however, Level (III) is not achieved, since the slopes of the fitted model and the bound model are significantly different.

## Appendix D: Polyominos: path complexity

Here we describe another approach to calculating complexity values for polyominos, which we call *path complexity*.

The path complexity method is based on ‘walking’ around the perimeter of the polyomino, and recording the direction of each step: F (forward), L (left), R (right). This yields a string of characters, which can be compressed. For example, a two-by-two polyomino made up of four tiles would have a short and repetitive path needed to describe its perimeter, and hence would have a low complexity value. By contrast, a larger and more irregular polyomino yielding a longer and more irregular path would be have a higher complexity value. Very few of the relatively small polyominos in the current data set have holes in them (i.e., missing tiles on the inside of the shape), so ‘walking’ around the perimeter of the shape is sufficient to describe the entire shape. For the few shapes which do have holes, we separately record the complexity of the hole and add it to the path complexity of the perimeter. In this manner, the path complexity corresponds to a descriptional complexity approach to assigning complexities, which is the essence of how Kolmogorov complexity quantifies complexity. (See also another ref. [152] for Kolmogorov complexity-based approach to estimating the complexity of polyominos).

In Figure D.2 this complexity measures works quite well, enabling the a priori prediction of the probabilities, just based on complexities of the shapes.

**FIG. D.2.**
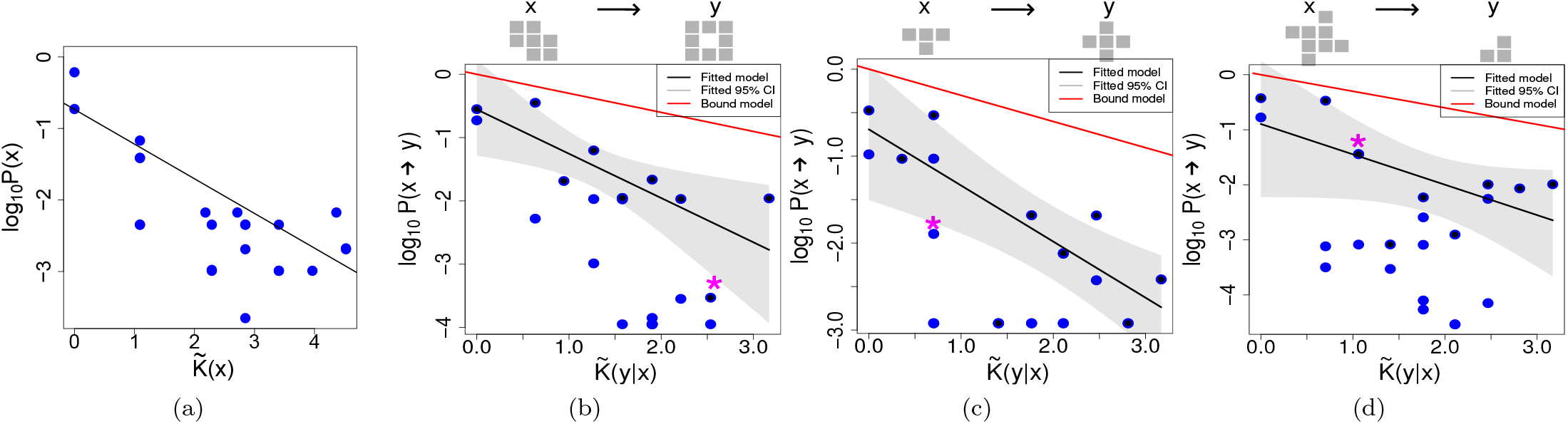
Self-assembling polyomino tiles (forward, left, right). (a) Simplicity bias found for the uniform random sampling of polyominos. The black line shows the linear regression using the maximal values for each category of complexity. On top of each plot we can see the phenotype x on the left side and an example of a phenotype y, into which phenotype x has mutated to. On the plots, the * indicates where the example y can be found. As previously explained, the following examples illustrate varying levels of success corresponding to different degrees of simplicity bias. (b) Example for the transition probabilities of a phenotype with complexity 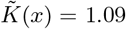. In this example *ρ* = − 0.57, such a negative correlation confirms Level (I). Level (II) is also achieved, with a *R*^2^ = 0.58. Finally, Level (III) is also achieved, since the differences of the slopes calculated with the bootstrap method and the slope of the bound model are not significantly different from 0. (c) In this example, using a phenotype of complexity 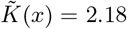, Level (I) is achieved with a *ρ* = − 0.42. Level (II) is also achieved, with a *R*^2^ = 0.59, however, Level (III) is also predicted, as slopes of the bound and fitted model are not significantly different. (d) In this example, using a phenotype of complexity 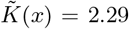, Level (I) is not achieved since the *p* − *value* = 0.18. Therefore, the rest of the levels are not achieved.

**FIG. D.3.**
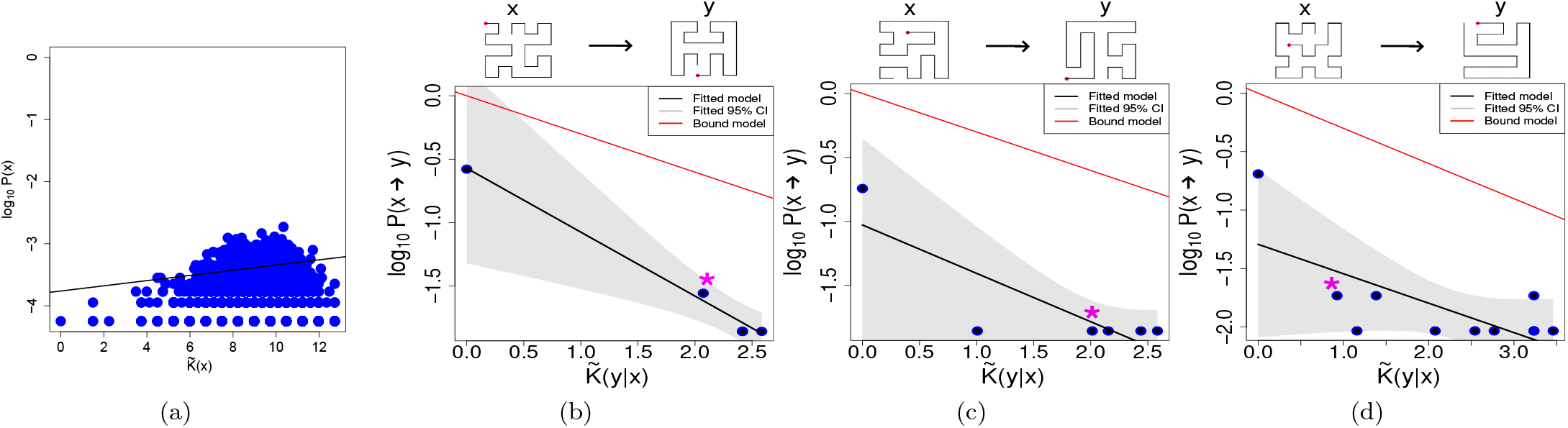
HP protein map compact L = 36. (a) Simplicity bias found for the uniform random sampling of HP maps. In the next plots, the black line shows the linear regression using the maximal values for each category of complexity. On top of each plot we can see the phenotype x on the left side and an example of a phenotype y, into which phenotype x has mutated to. On the plots, the * indicates where the example y can be found. As previously explained, the following examples illustrate varying levels of success corresponding to different degrees of simplicity bias. (b) Example for the transition probabilities of a phenotype with complexity 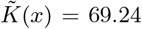. In this example *ρ* = − 0.89, such a negative correlation confirms Level (I). Level (II) is also achieved, with a *R*^2^ = 0.99. Finally, Level (III) is also achieved, since the differences of the slopes calculated with the bootstrap method and the slope of the bound model are not significantly different from 0. (c) In this example, using a phenotype of complexity 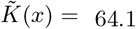, Level (I) is not achieved, since the *p* − *value* = 0.16. Next levels are therefore also not achieved. (d) In this example, using a phenotype of complexity 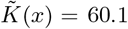, Level (I) is not achieved since the *p* − *value* = 0.09. Therefore, the rest of the levels are not achieved. The levels of success for the different levels is, considering genotypes Level I: 0.09; Level II: 0.09; Level III: 0.87. Considering phenotypes: Level I: 0.08; Level II: 0.07; Level III: 0.85. As before, these proportions of success are based on the total number of genotypes and phenotypes predicted in the previous level.

## Appendix E: Conditional complexity for found vs. not found phenotypes

For any given starting phenotype *x*, random mutations to its underlying genotypes may or may not yield all possible other phenotypes. In fact, it is quite likely that only a reduced set of phenotypes are accessible via single-point mutations from a given starting phenotype *x*, in other words *P* (*x* → *y*) = 0 for most *y* in the set of all possible phenotypes. We might expect that those phenotypes with lower conditional complexity will be ‘found’ via point mutations, where found ‘means’ that *P* (*x* → *y*) *>* 0, or in other words the phenotype *y* is present in the uniform random sampling and in the 1-mutational neighborhood of a given phenotype *x*. Similarly, we would conjecture that those with higher conditional complexity will not be found. The intuition here is that the conditional simplicity bias bound gives higher *a priori* probability to similar (or rather, lower conditional complexity) phenotypes, and so presumably the ‘not-found’ phenotypes will be those of highest conditional complexity.

We test this conjecture for the maps studied above by calculating the conditional complexity 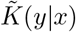 of all the found phenotypes and the conditional complexity 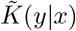 of the phenotypes that were not found. Next, we calculate the median conditional complexity for both groups and determine the difference between them: 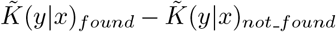. This process is repeated for each explored starting phenotype *x*. To facilitate comparison between the different models, we normalize each group by dividing by the absolute maximum value within that group. According to the conjecture stated above, we expect that 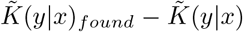 _*not found*_ is negative (or at least not positive). In Figure E.4 the results of the numerical experiments are shown. Most cases display a negative value of 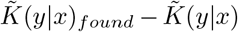 _*not found*_, and for none of them it is positive, in line with our expectation, as most of the found phenotypes have a lower conditional complexity than those not found, with only a few exceptions.

In this analysis, it is important to note that, aside from the polyominos and the HP protein map, the sampling process is incomplete. Specifically, we are unable to identify all possible phenotypes or fully explore the entire 1-mutational neighborhood for each of the studied phenotypes, as both genotypes and phenotypes are continuous. A more comprehensive exploration could potentially uncover very rare phenotypes, which might influence the results presented here. However, these rare phenotypes are likely to be more complex than those identified in our current exploration. Under such circumstances, our conjecture would likely become even more apparent in this analysis.

**FIG. E.4.**
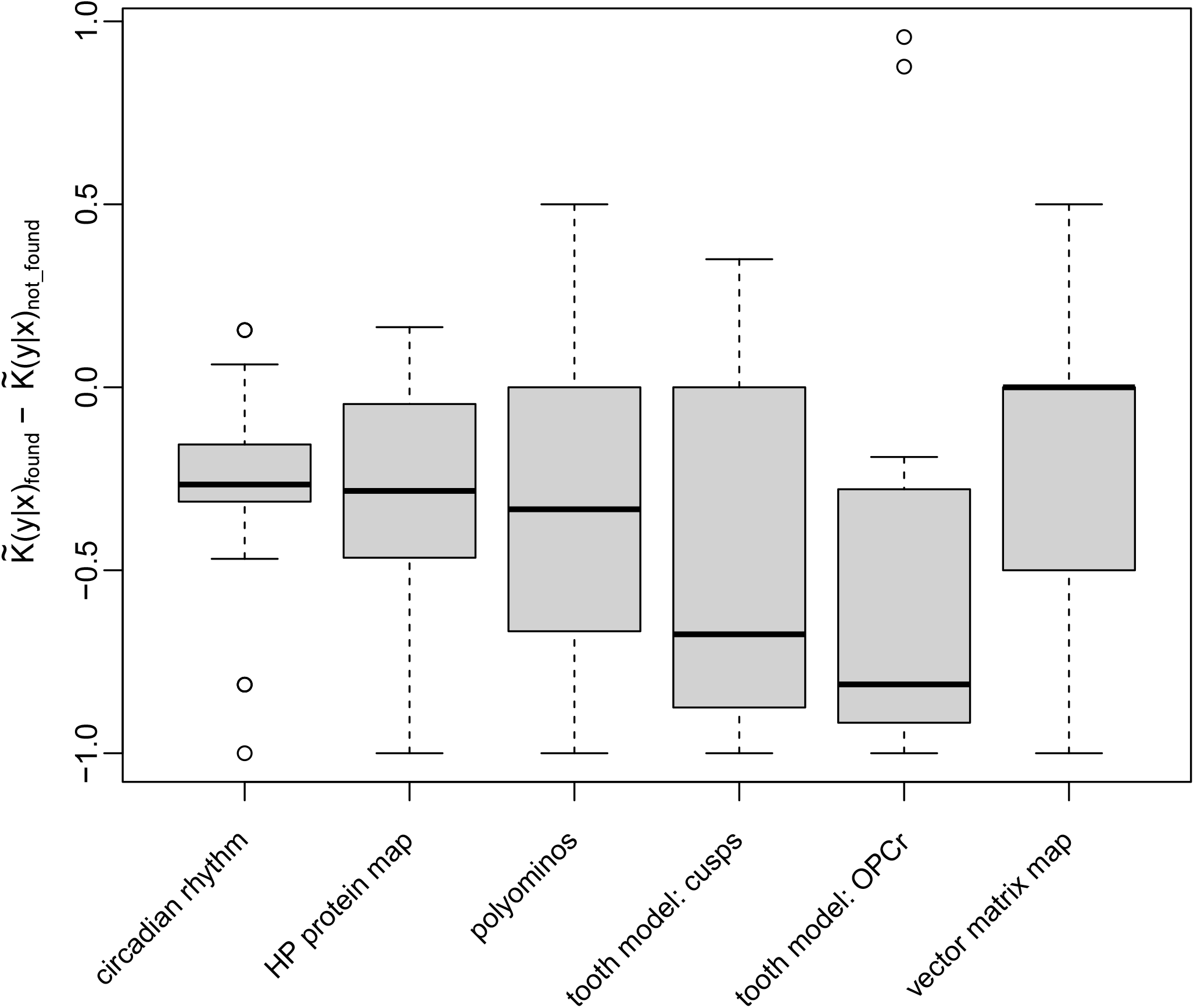
Phenotypes that are more frequently found tend to have lower conditional complexity. This figure shows the difference between the median conditional complexity of “found” and “not found” phenotypes, normalized by their maximal value. A “found” phenotype is one identified both through uniform random sampling and through single-point mutations of a phenotype *x*, such that *P* (*x → y*) *>* 0, while a “not found” phenotype is one observed only in uniform random sampling and not through single-point mutations, meaning *P* (*x → y*) = 0. The median conditional complexity is calculated for both groups, and for each phenotype *x*, the difference between these medians is determined as 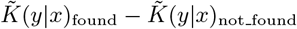. Negative values indicate that the conditional complexity 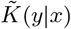 of the “not found” phenotypes is higher. The boxplots present the results of these calculations for various systems, with the number of phenotypes *x* explored specified for each: Circadian rhythm (*n* = 35), vector matrix map (*n* = 310), polyominos (*n* = 21), HP protein map (*n* = 26), teeth model: OPCR (*n* = 12), and teeth model: cusps (*n* = 12).

